# Dynamic domain interactions encode possible CheA autophosphorylation mechanisms revealed by coarse-grained simulations

**DOI:** 10.64898/2025.12.21.695815

**Authors:** Jian Huang, Katherine Wahlbeck Lu-Diaz, Lynmarie Thompson, Jianhan Chen

## Abstract

Autophosphorylation of CheA is key to initiation of the phosphorylation cascade that eventually controls the direction of downstream flagellar motors for chemotaxis signaling in motile bacteria. The phospho-transfer reaction, from ATP bound in the P4 catalytic domain to a specific His residue in the P1 substrate domain in CheA, can be significantly accelerated within core signaling unit complexes containing chemoreceptors, CheA and CheW. Previous studies have proposed that CheA autophosphorylation activity is regulated by changing the dynamics of P4 and/or altering its interactions with P1 in response to signals transmitted from chemoreceptors. However, the positions of CheA P1 and P2 domains in the core signaling unit are not well-characterized because they form only transient interactions with other domains. Though previous studies have identified possible domain-domain interaction surfaces, especially for P1 and P4, a bottom-up analysis of these interactions has not been performed. Here, we employed extensive molecular simulations to analyze interactions among CheA domains using a hybrid resolution (HyRes) protein model designed for dynamic protein structures and interactions. The results revealed multiple major modes of dynamic P1/P4 interactions. In particular, P1 was found to bind dynamically near the preferential binding surfaces on P4 even in the ATP free state. ATP binding to P4 reduces the motion of P1 binding and promotes a *trans-* productive-like mode that brings His48 in close contact with the bound ATP for possible autophosphorylation. The predicted non-productive and productive P1/P4 interaction modes appear highly consistent with existing NMR, mutagenesis, and chemical modification data. Together, these findings provide a more complete picture of the dynamics of domain-domain interactions of CheA and new insights into the possible regulation mechanism of its autophosphorylation.

## Introduction

The chemotaxis signaling pathway is a critical mechanism that various motile bacteria employ to detect chemical gradients of environmental chemoeffectors and swim towards attractants or away from repellents (1–4). Chemotaxis signaling begins with sensing by highly ordered supramolecular arrays of core signaling units (CSUs) on the membrane (5, 6). The CSU, the minimal unit of chemosensory function, consists of the transmembrane receptors known as methyl-accepting chemotaxis proteins (MCPs), a histidine autokinase CheA, and a coupling protein CheW (5, 6). Chemosensory arrays formed by assembled CSUs enable cooperative signaling of the MCP receptors upon small changes of chemoeffector concentration that cause large changes of the autophosphorylation activity of the CheA autokinase (7–9). Once CheA phosphorylates itself, it can transfer its phosphate group to downstream response regulators, including CheY, which interacts with the flagellar basal body to drive changes in the direction of motor rotation of the flagella that biases random walks towards favorable environments (1).

**Figure 1.**
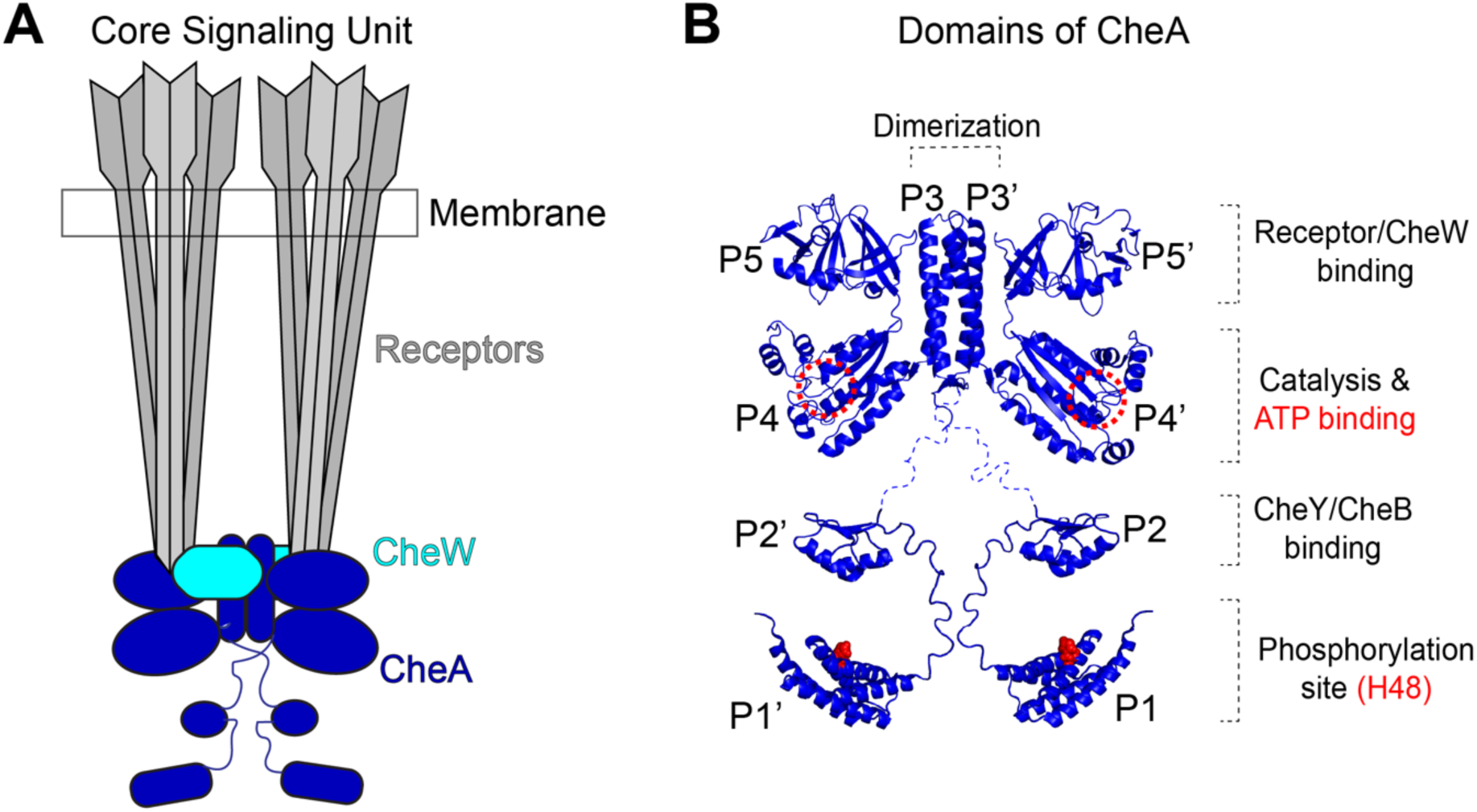
Core signaling unit in chemotaxis and the autokinase CheA. **A)** Schematic topology of the core signaling unit for chemotaxis with key components annotated. **B)** CheA autokinase dimer structure with all five domains and some key functions or features annotated (PDB: 1B3Q (10) used for domains P1-P2 and PDB: 2LP4 (11) used for P3-P4-P5).

By fitting electronic densities from cryogenic electron tomography (cryo-ET) with available experimental or AlphaFold2 predicted structures, several models of the overall molecular architecture of the *E. coli* CSU complex have been constructed (12–14). In the CSU complex, the membrane-bound MCP receptors are organized as two trimers of dimers (TOD), which are bridged by a CheA homodimer and bound with two CheW monomers (13–16). Each CheA monomer has five domains, P1-P5, connected by flexible linkers or short hinge regions (17). The P3/P3’ in the CheA dimer forms a four-helix bundle as the principal dimerization element, and is positioned between the two receptor TODs (12, 16, 18). The CheA.P4 domain harbors an ATP-binding pocket and catalyzes the *trans-* phosphorylation of H48 in the P1’ domain of the other monomer (19, 20). The P2 domain interacts with the response regulators (CheB and CheY) for downstream signal transduction or adaptation modulation (21, 22). Even though P3, P4 and P5 domains are clearly discernable and fit well to the electron densities in the current cryo-ET studies (12, 16, 18), the P1 and P2 domains cannot be unambiguously modeled into the remaining amorphous electron density, likely due to their dynamic interactions with the rest of the assembly.

Previous studies have shown that the kinase activity changes of CheA do not result from large scale rearrangements of the order or packing of chemosensory arrays upon binding of chemoeffectors (23–28). Instead, current evidence suggests that smaller scale changes such as changing of CheA domain positions or stabilities controls the kinase activity and that ATP binding is essential for the P1/P4 interactions required for autophosphorylation (28, 29). Studying the dynamic domain-domain interactions in CheA is thus important to understand the catalytic and regulatory mechanisms of its kinase activity. Several studies have reported on P1/P4 interactions in CheA and have identified the binding surfaces on both P1 and P4 for these interactions. An early chemical shift perturbation NMR study of CheA domains identified a non-productive binding mode, where the ATP binding site on P4 and the substrate histidine of P1 are distant and autophosphorylation could not occur in this interaction mode (30). The P1/P4 binding surface was suggested to involve residues D93, M94, R97 (*T. maritima* numbering) in helix-D of P1, and E387 and V396 of P4, which are located on the opposite side of P4 to the catalytic ATP-binding pocket (30). This non-productive binding mode is proposed to be favored in the inactive CheA state since it prevents close contacts between ATP and the P1 substrate histidine.

A putative productive mode involving helices A and B of P1 and residues around the ATP binding site of P4 was first produced through protein-protein docking followed by short 4-ns molecular dynamics (MD) simulations (31). The essence of the predicted interaction interface on P1 appears consistent with a later mutational analysis, which identified a set of surface-exposed residues distal to His-48 (32). The mutational screening of P1 confirmed the involvement of helices A and B by reporting a series of sites on helix-A and the start of helix-B of *Escherichia coli* P1 (T11, D14, E18, H26, E38 and A42), where mutations decrease the phosphorylation rate (32). A systematic cysteine substitution and bulky probe attachment study pinpointed P4 residues S351, D359, I388, R466, and V485 of the *Salmonella Typhimurium* CheA P4 as the possible P1 domain binding site (33). More recently, Muok *et al.* revealed several interaction pairs between P1 and P4 domains from *T. maritima* using chemical cross linking followed by mass spectrometry (34). Kinetic measurements suggest an ordered sequential mechanism, with ATP binding before P1. The productive interaction mode was proposed to be dependent on the presence of ATP in the P4 binding pocket, which was proposed to induce a reordering of the ATP lid loop of P4 (35) as observed in one nucleotide-bound crystal structure of P4 (36), or an opening of the ATP lid to allow access for P1 binding (31). Thus, the ATP lid loop conformational change could contribute to the formation of the productive P1/P4 interface.

P1 domains may dimerize within CheA dimers in addition to non-productive and productive interaction modes between P1/P4. P1 domains in the CheA dimer have been shown to interfere with each other during the *trans-* autophosphorylation reaction, because the reaction rate increases when one P1 domain subunit is deleted (37, 38). A P1 dimer model was previously reported based on distance restraints derived from pulse-dipolar electron-spin resonance (PDS) spectroscopy measurements (34). In this model, based on crystal contacts in one of the P1 monomer structures (PDB: 1TQG (39)), the two P1 domains are packed in a parallel fashion using the helix-A and helix-B as the dimerization interface where the two H48 substrate sites face towards each other (34). Tran *et al* (29) and Cassidy *et al* (18) constructed antiparallel models of P1 dimers with helices A and D or helices A and B at the dimerization interface, respectively. If P1 dimerizes to a significant extent using helices A and B as the dimerization interface, it could inhibit the autophosphorylation reaction since H48 sidechains are occluded. Thus it is important to determine the type and extent of P1/P1’ interactions.

Physics-based MD simulations are arguably necessary to overcome the challenges of obtaining a high-resolution molecular-level description of the dynamic domain-domain interactions of the CheA dimer. However, the large size of the CheA dimer and long conformational exchange timescale are inaccessible to conventional all-atom explicit solvent simulations (40, 41). By reducing all-atom resolution to a lower level, coarse grained (CG) models can effectively decrease degrees of freedom, facilitate the sampling of protein conformations, and greatly expand the accessible timescales (42, 43). Over the past few decades, CG models have shown powerful applications in structure prediction, protein folding, as well as protein-protein interactions (42, 43). In particular, a Hybrid resolution (HyRes) CG protein model has been recently developed for accurate description of dynamic protein conformations and interactions (44, 45). In HyRes, the backbone is represented at all-atom level, to capture backbone-mediated interactions and secondary structures, and side chains are represented at intermediate resolution with up to 5 beads, allowing semi-quantitative description of transient long-range interactions. It has been demonstrated that HyRes provides an accurate description of residual helicity profiles of over a dozen intrinsically disordered proteins (IDPs), captures the radius of gyration (*R*_g_) of ∼30 IDPs with ∼0.7 correlation (46), and recapitulates NMR PRE profiles of p53-TAD at a level comparable to the latest a99SB-disp all-atom force field (47). HyRes has proven highly effective in simulating dynamic protein interactions including phase transitions (46–49), making it uniquely suitable for studying conformational dynamics of the CheA dimer.

Herein, we deployed HyRes to study the dynamic domain-domain interactions of the CheA dimer in the absence and presence of ATP bound in P4. Comparison of the domain placement distributions within the dimer revealed that P1 has inherent propensity to interact dynamically with P4, resembling the non-productive modes identified in previous studies. With ATP bound to P4, HyRes simulations successfully sampled closer contacts between P1 and P4, yielding productive-like binding configurations that agree well with existing experimental data. Finally, the most likely transient P1/P1’ dimer interactions were found to occlude H48, explaining the ability of P1/P1’ interactions to interfere with autophosphorylation. Results from this bottom-up simulation study provide valuable insights into the dynamics of domain interactions of the CheA dimer and possible regulatory mechanisms for autophosphorylation.

## Methods

### AlphaFold2 modeling of the *E. coli* CheA dimer structure

The UNIPROT accession number (P07363) sequence isoform 1 was used for the AlphaFold2 mmseqs2 prediction (https://github.com/sokrypton/ColabFold) to generate five structure models. These models are highly similar in the P3-P5 domains, while the P1 and P2 domains show modest differences (Figure S1). All models were then relaxed using the Rosetta fastrelax protocol (50, 51) with the “constrain_relax_to_start_coords” flag on and the “ramp_constraints” flag on to avoid large deviations from the original coordinates. The relaxed structure with the lowest Rosetta energy (model 4) was used as the starting point for further modeling and simulations.

### HyRes-compatible CG modeling of ATP

ATP is represented using the MARTINI coarse-grained mapping (52) due to the similarity to HyRes mapping of protein side chains (see Figure S2A). Force field parameters for bonds and dihedrals such as the equilibrium bond lengths and force constants, and dihedral angles and force constants are directly adopted from the MARTINI ATP model. The equilibrate values of all angles were also adopted from the MARTINI ATP, while the force constants of angles were adjusted by scaling down the original values by a factor of 20, to match the level of covalent flexibility in HyRes. For non-bonded parameters, we used the same *R*_min_ as in the MARTINI ATP whereas the epsilon values were first adopted from the MARTINI interaction table (53) and then were scaled down by a factor of 33. This factor was to account for implicit treatment of solvent in HyRes, and was determined by the averaging scales of epsilon values of Leu, Phe and Trp residues between MARTINI and HyRes. Four improper dihedrals were set up to maintain the planar structure of the adenosine ring. The CHARMM format of the parameter and topology of the HyRes ATP is provided in the supplementary information. The conformation of ATP bound to the CheA dimer was derived from the previously resolved ATP analog-P3P4 structure from *Thermotoga maritima* (PDB: 4XIV) (37). Note that the bound ATP was positionally restrained during all simulations in this work; the exact choices of covalent and vdW parameters are not expected to be important in studies of CheA dimer conformational dynamics, except for the long-range electrostatic force.

### HyRes simulation of the CheA dimer

The Rosetta-relaxed model (Figure S1, model 4) was converted to the HyRes CG representation using “at2HyRes” python scripts (https://github.com/mdlab-um/HyRes_GPU). HyRes is designed for simulation of dynamic protein conformations and folded domains require internal restraints to maintain the tertiary structure. For P1, P2 and P4, all pairs of Cα atoms within 12 Å from regions with secondary structures were subjected to distance restraints with a small force constant of 100 kJ/mol/nm^2^. Considering that the P3 and P5 domains are tightly packed with the CheW and the cytosolic part of the chemotaxis receptors in the signaling complexes, we positionally restrained all Cα atoms of these two domains with a modest force constant of 400 kJ/mol/nm^2^ using the initial structure as the reference. To generate randomized initial positions of the P1 and P2 domains in the dimer, high temperature simulations were performed at 500K for 200 ns in OpenMM8 (54), using the LangevinMiddleIntegrator with a friction of 0.1 per ps. The non-bonded interactions were gradually switched to 0 from16 Å to 18 Å. The salt concentration was set at 0.15 M and an effective dielectric constant of 20 was used. The Verlet integrator with an integration timestep of 2 fs was used, and snapshots were saved every 200 ps. From the resulting trajectory, six snapshots were randomly selected and used as the initial configurations. Each of these configurations was used to initiate an independent 8-µs simulation at 300 K using the same MD configuration except with an integration timestep of 4 fs. The convergence of these simulations was assessed by splitting each trajectory into the first and second halves and comparing calculated properties from all combined first and second halves (e.g., see Figure S3).

For the ATP-bound state, distance restraints with the contacting residues in the binding site of P4 were imposed with a force constant of 400 kJ/mol/nm^2^ (Figure S2B). The distance restraints were derived from a previously reported ATP analog-P4 kinase domain complex structure from the *Thermotoga maritima* (36). The ATP-coordinating residues are assumed to be conserved based on the sequence homology and are mapped to the sequence of the P4 kinase domain of the *Escherichia coli* using sequence alignment (Table S1). Specifically, the distance restraints include: between D420 Cα and the center of mass (COM) of the adenine group of ATP (NA, NB, NC and ND beads) with a distance of 10 Å, between H376 Cα and the COM of the three phosphate beads (P1, P2 and P3) of ATP with a distance of 8 Å, R379 Cα and the COM of the three phosphate beads of ATP with a distance of 8 Å. A set of 6 independent simulations were performed following the same protocol as outlined above for the ATP-free state.

### Simulation of CheA.P1 dimerization

To examine possible P1/P1’ dimer configurations, the CheA.P1 structure was extracted from the AlphaFold2 predicted model-4, and then used to create a random initial configuration with a large COM separation of 50 Å using the packmol software (55), which was then converted to the HyRes representation. A flat-bottom restraint with a force constant of 400 kJ/mol/nm^2^ was used to keep the COM of the two CheA.P1 within 50 Å. Each CheA.P1 was also internally restrained to maintain the folded structure using the same procedure as described above. The system was simulated using the HyRes force field for 2 µs after a short 100 ps equilibration. The same MD configuration as above was used except for an elevated temperature of 380 K, which help accelerate the sampling of various possible dimer configurations. Because each monomer is internally restrained, the elevated temperature should not affect the relative abundance of various dimer states.

### Domain-domain contact analysis

Analysis of domain-domain contacts and distance distributions for both the ATP-bound and ATP-free systems were performed using MDAnalysis with in-house scripts (56). The first 500 ns of all trajectories were removed when conducting contact analyses. To analyze *trans-* productive contacts of P1/P4, distance calculations were performed either between H48 and the terminal phosphate bead of ATP or R379 (an ATP-coordinating residue in the bound state of P4). For the proposed non-productive P1/P4 binding mode (30), we first mapped those P1 and P4 residues that showed chemical shift perturbations in the prior study of *T. maritima* CheA to the *E. coli* CheA based on sequence alignment (Table S1), and then calculated distances between the Cα atoms of S358 and I367 of P4 to the COMs of D96, I97 and E100 on P1. To characterize the P1 domain COM density distributions, the COMs of P1 domains from all trajectories were first calculated and then combined to derive the DX volumetric density maps for visualization in VMD (57). To convert the HyRes representations to the all-atom ones, we extracted the atomistic backbone coordinates and used the pdbreader tool from CHARMM-GUI to rebuild the sidechains (58, 59).

To analyze the contacting mode between the P1 and P4 domains, we first extracted the snapshots forming the *trans-* productive contacts between P1’ and P4 or P1 and P4’ using a distance cutoff of 8 Å between the Cα atom of P1.H48 and the terminal phosphate group from the P4.ATP. These snapshots were characterized using the distance between the COMs of the helix-A (residue 9-27) of the P1 domain and the α2 helix (residue 369-385) of the P4 domain, and the angle between the helix-A of the P1 domain (defined as a vector pointing from the C-α atom of residue 14 to the C-α atom of residue 25) and the α2 helix of the P4 domain (defined as a vector pointing from the C-α atom of residue 370 to the C-α atom of residue 381). Regions with the highest densities were identified as the major binding modes.

To analyze the contacting mode from the P1/P1’ dimerization simulations, we analyzed all snapshots sampled during the 2 µs simulation at 380 K, using the interhelix COM distance and interhelix angle between the two helix-B from the two P1 domains (residue 38-55 with residue 41 and 52 define the directional vector). Representative atomistic models of the P1/P1’ dimer were generated from HyRes structures using the pdbreader in CHARMM-GUI (58, 59).

## Results

### Generation of starting structures of full length CheA for HyRes simulations

To obtain starting structures of the CheA dimer for HyRes simulation, we predicted the full-length *E. coli* CheA dimer structure using AlphaFold2 and refined them with Rosetta (see Methods). As shown in Figure S1, the five AlphaFold2 models of the CheA dimer show consistent tertiary structures for all domains as well as the relative positioning of P3-P5 domains, even though linkers connecting P1/P2 and P2/P3 vary significantly and have large variability and lower pLDDT values (Figure S1). The P2 domains have minimal contacts with other domains in all models. Those features are consistent with the model previously derived by Cassidy *et al* based on cryo-electron tomography and molecular simulation (18), with a small Cα RMSD of ∼3 Å for P3-P4-P5 domains. Similar to Cassidy *et al*’s model (18), the P1/P1’ domains are predicted as an antiparallel dimer with the phosphorylation site H48 shielded and away from the active site on P4 domains in a non-productive P1/P4 binding mode. However, the 5 AlphaFold2 structures differ from each other and Cassidy *et al*’s model with slightly different P1/P1’ packing angles (Figure S1). The AlphaFold2 models clearly do not capture the presumably dynamic interactions that P1 must have with P4 for autophosphorylation to occur.

### CheA P1 domain binds dynamically to P4 domains in an ATP-dependent fashion

To probe the domain-domain interactions in the CheA dimer, especially between the substrate P1 domain and the kinase P4 domain, six independent 8-µs HyRes simulations were performed with randomized initial positioning of the P1 and P2 domains for both the ATP-bound and ATP-free states (see Methods). The simulations reveal large-scale dynamics in the position of P1 and P2 domains and their interactions with the rest of the CheA dimer (e.g., see SI Movies S1 and S2). Block analysis (see Methods) demonstrated that the COM densities of P1 and P2 domains derived from the first and second halves of the simulations were in general agreement (Figure S3), indicating good convergence. In HyRes simulations, we observed that there are two major locations of the P1 and P2 domains with respect to the stable P3-P4-P5 domains, with a docking region near the ATP binding pocket (termed the “upper” docking region, Figure 2A middle panel) and another docking region directly below the P4 domain (termed the “lower” docking region, Figure 2A middle panel). The predicted “lower” docking regions are reminiscent of the unresolved densities observed in cryo-ET experiments (12, 18, 28). It is noteworthy that, even without ATP bound to P4, P1 domains show a significant propensity to interact with P4 near the ATP binding site, in both *cis-* (P1/P4) and *trans-* (P1/P4’) configurations (Figure 2). Strikingly, in the presence of bound ATP, the *cis-* P1/P4 “upper” docking region was largely eliminated (magenta meshes in Figure 2 left column and blue meshes in Figure S4). In contrast, the P2 COM densities are almost the same with and without ATP bound (cyan and grey meshes in Figure 2 right column). This is an interesting prediction supporting that ATP binding dramatically modulates the P1 and P4 domain interactions to promote *trans-* and likely productive contacts for autophosphorylation, as suggested experimentally (28, 29).

**Figure 2.**
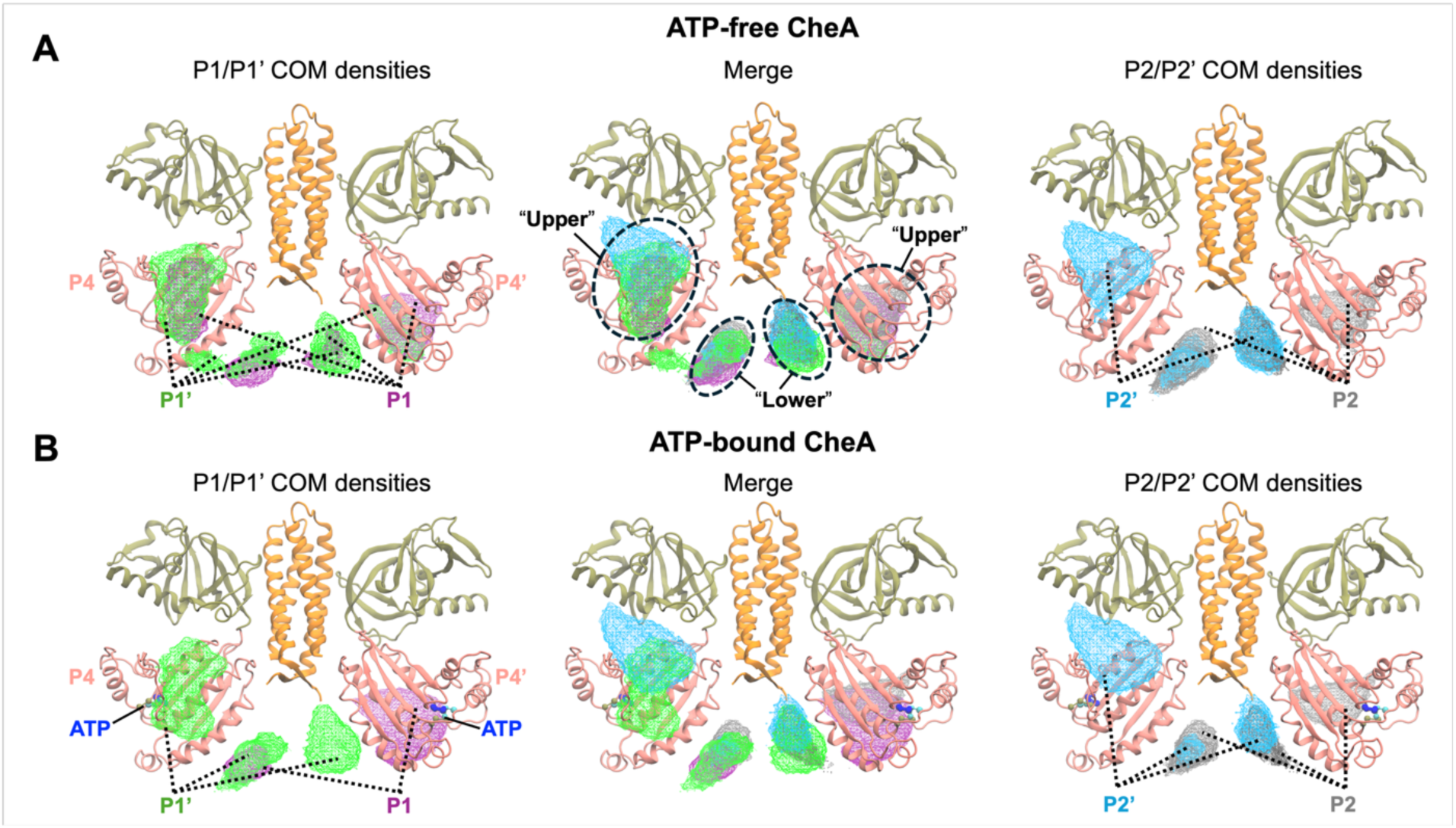
Distribution of P1 and P2 COM in the ATP-free (A) and ATP-bound (B) states of the CheA dimer. Only the P3, P4 and P5 domains are shown in cartoon representation, while P1/P2 domains and linkers are hidden for clarity. The COM density maps for the P1 and P1’ domains are shown as magenta and green meshes, and those for the P2 and P2’ domains are shown as grey and cyan meshes. All COM density maps are shown using an iso-value of 12 e/Å^3^. The bound ATP is shown as ball-and-stick with beads forming the adenine group, the ribose ring, and phosphate groups colored in blue, cyan, and olive, respectively. The P1/P1’ densities and P2/P2’ densities are shown separately in the left and right panels, with the merged ones shown in the middle panels.

### Bound ATP promotes close *trans-* contacts between P1 and P4 domains

To further evaluate whether ATP binding promotes close *trans-* P1/P4 interactions that could enable autophosphorylation, we analyzed all P1/P4 configurations in the “upper” docking region in both states by using the interhelix distance and angle between the helix-A of P1 and the helix-α2 of P4 (see Methods). As shown in Figure 3A, although ATP did not significantly change the locations of minima of favored P1/P4 interacting configurations, it did shift the relative density of their occupancies to shorter interhelix distances (< 25 Å). To check if these differences in P1 interaction are related to an increase in productive binding, we further analyzed the distance between H48 (phosphorylation site) on the P1 domain and bound ATP on the P4 domain in the “upper” docking regions. Figure 3B&C show the distributions of distances between the H48 Cα atom and the terminal phosphate group of ATP molecules (in the ATP-bound system) or the ATP-coordinating R379 residue (in the ATP-free system). R379 has been shown to coordinate the terminal phosphate group of ATP in the previously determined structure (36) and thus was chosen as a good approximation for the distance calculation in the ATP-free system. As summarized in Figure 3C, in the ATP-free state, *trans-* P1/P4 interactions rarely sample any conformation with H48 and R379 within 15 Å (green shaded area). In contrast, close *trans-* H48-ATP contacts were consistently sampled in simulations of the ATP-bound state, in five out of the six trajectories (Figure 3C). That is, in addition to eliminating the *cis-* P1/P4 “upper” interaction, the presence of ATP molecules in P4 domains modulates the docking configuration of the P1 domain such that H48 has a significantly enhanced probability of making close *trans-* productive interactions with the ATP phosphate group, potentiating the autophosphorylation process. Note that using the same R379 to H48 Cα in the ATP-bound simulations gives a similar picture of close contact formation (Figure S5). Also note that the ATP-lid was modeled as a disordered loop in the initial configurations of all systems instead of an ordered ATP-lid as seen in one crystal structure of the P4 domain bound to a nucleotide and Mg^2+^ (36). It is possible that the disorder-to-order transitions of the lid could further promote the productive-like *trans-* contacts between the P1 and P4 domains. Nonetheless, the apparent ability of HyRes simulations to recapitulate the ability of ATP binding to promote productive *trans-* P1/P4’ contacts is noteworthy.

**Figure 3.**
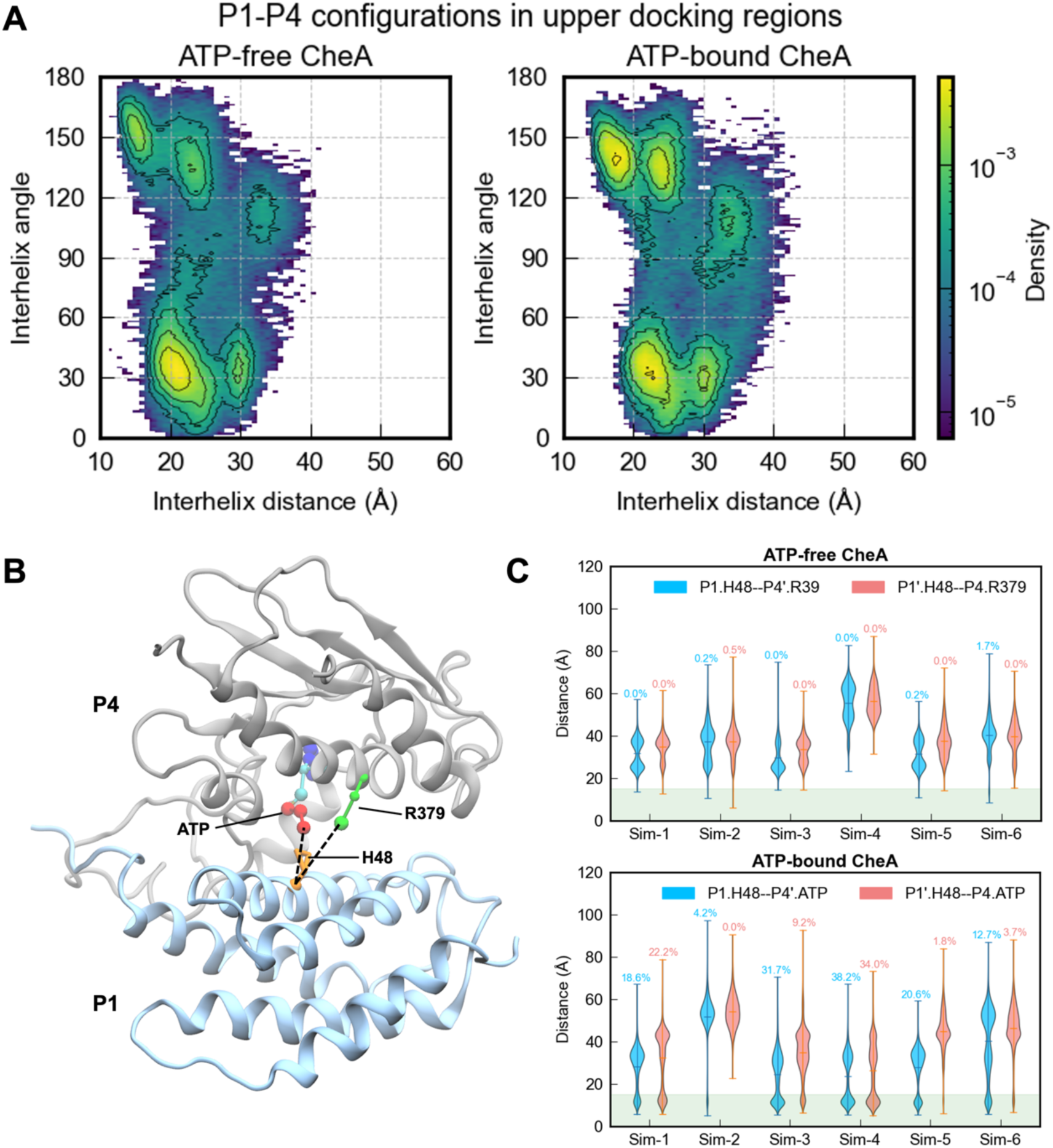
Analysis of close *trans-* P1/P4 interactions. **A)** Probability distributions of P1/P4 configurations in the “upper” docking regions in ATP-free and -bound states as a function of the interhelix distances and angles between the helix-A of P1 and the helix-α2 of P4 (see Methods). **B)** A representative snapshot of P1 and P4 forming close *trans-* interactions potentially capable of autophosphorylation observed in simulations of the ATP-bound state. The P1’ and P4 domains are shown as gray- and light-blue-colored cartoons. R379 and H48 (in HyRes resolution) are shown by green and orange sticks. The bound ATP is shown as balls and sticks with beads forming the adenine group, the ribose ring and phosphate groups colored blue, cyan and red, respectively. C) Violin plots of distance probability distributions from six parallel simulations of the ATP-free (upper panel) and ATP-bound state (lower panel). Note that the distance from P1.H48 Cα to P4’.R379CC bead (the tip) was calculated for the ATP-free state and to ATP.P3 bead bound in P4’ was calculated in the ATP-bound state. Also note that violin plots are similar for the ATP-bound system measured to R379, as shown in Figure S5. The two *trans-* distances (P1.H48 to P4’.ATP or P4’.R379 and P1’.H48 to P4.ATP or P4.R379) are plotted separately with cyan and salmon colors for each simulation trajectory. The green shade area marks contact distance within 15 Å, with the probabilities of forming < 15 Å labelled above each violin plot.

### Predicted productive-like *trans-* P1/P4 interactions are consistent with available experiments

We further characterized the predicted *trans-* P1/P4 productive binding mode and interaction interfaces in the ATP-bound state. For this, we selected snapshots using a distance cutoff of 8 Å between the H48 Cα (H48.Cα) and the terminal phosphate of ATP (ATP.P3) to include only the most tightly bound P1/P4 interactions. The cutoff distance was chosen to reflect the sum of the average distance between H48’s Cα and either nitrogen atom of its imidazole is (∼ 3.5 - 4.5 Å) and the size of the phosphate group of ATP (radius is about 2.2 Å). As such, the selected configurations better represent a productive-like state that could support autophosphorylation. As shown by the red spheres in Figure 4A, the COMs of P1’ domain from these productive frames are clustered within a small subsection of the “upper” docking region. Although the P1 COM of the ATP-free state also samples this region (Figure 2A), it does not adopt the orientation needed for making close H48-ATP contacts (Figure 3C).

**Figure 4.**
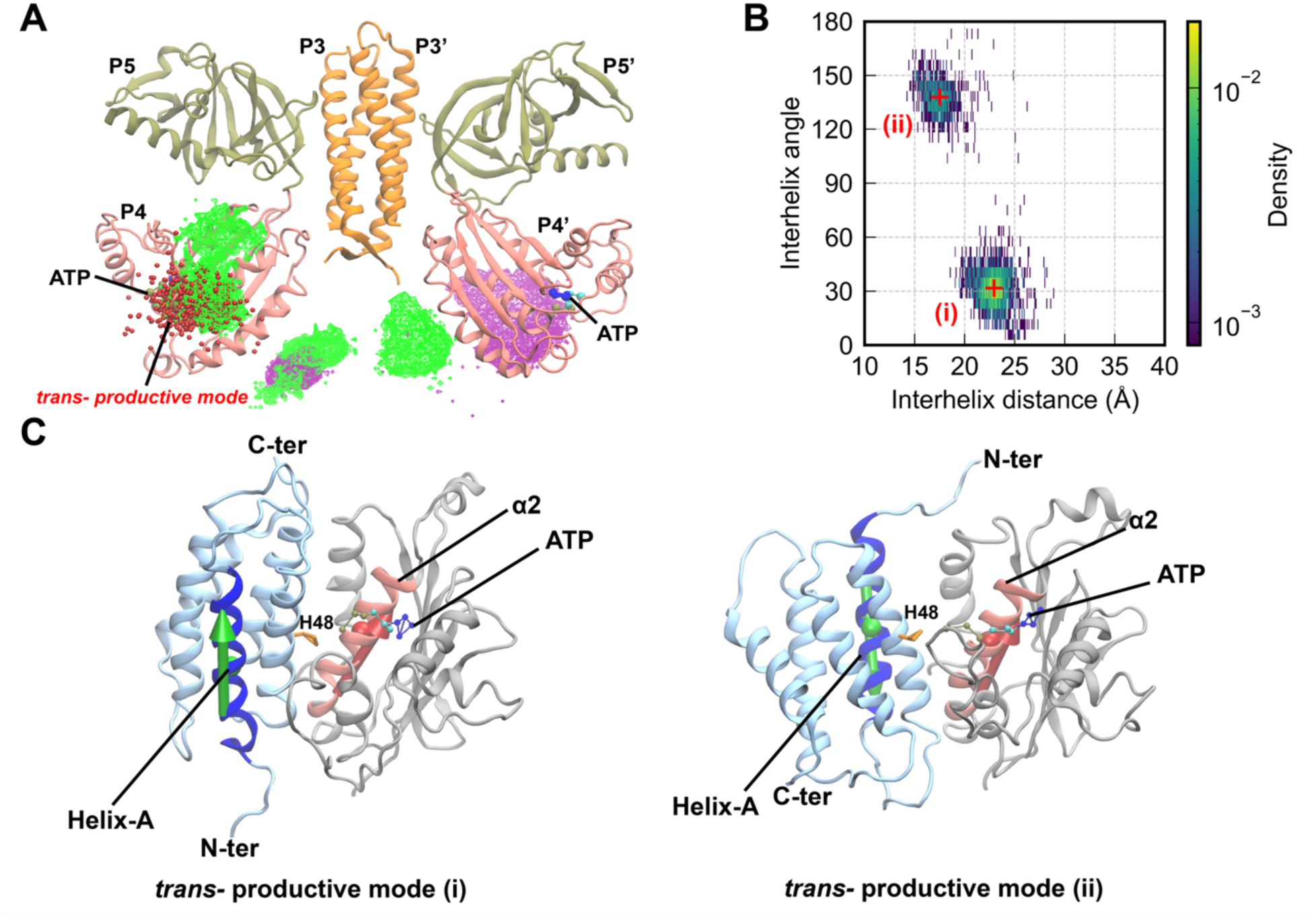
P1/P4 domain binding modes from all *trans-* productive snapshots. **A)** COMs of the P1’ domains for P1/P4 productive binding (with Cα of H48 within 8 Å of the ATP terminal phosphate) are plotted using red spheres along with the full P1 and P1’ COM densities from the ATP-bound CheA system (green and purple meshes), using the same coloring and labeling scheme as in Figure 2A. **B)** Probability distribution of the productive-like *trans-* binding configurations as a function of the COM distance and directional vector angle between helix-A of the P1 (or P1’) domain and helix-α2 of the P4’ (or P4) domain (see panel C). **C)** Two representative snapshots of the major P1/P4 binding modes, corresponding to the cluster center (i) and (ii) (marked by red “+”). The directional vectors of the helix-A of P1 and helix-α2 of P4 are shown as green and red arrows, respectively. The COMs of the helix-A and the helix-α2 helix are shown as green and red spheres. The substrate H48 is shown as orange sticks, while the ATP molecule is shown as ball and sticks with beads forming the adenine group, the ribose ring and phosphate groups colored blue, cyan and red respectively.

To determine if the predicted productive-like binding interactions are consistent with existing experimental data on P1/P4 interactions or the previously predicted Zhang’s productive model (31), we first characterized the productive frames of simulations using the interhelix COM distance and the cross angle between P1 helix-A and P4 helix-α2 (see Methods; Figure 4C). There are apparently two major modes of productive *trans-* binding (Figure 4B). The first mode, labeled as (i), accounts for ∼ 70% of all productive frames and has an interhelix COM distance of 23 Å and a directional vector angle of 31.7°; the second mode, labelled as (ii), accounts for ∼ 30% of all productive frames and has a COM distance of 17.5 Å and a directional vector angle of 138°. P1 in the mode (i) packed nearly in a parallel orientation, while P1 in mode (ii) adopted an antiparallel orientation of P1 relative to P4 (see green arrows on P1 with respect to pink arrow on P4 in Figure 4C).

The HyRes-predicted *trans*-productive modes are consistent with experimental evidence for P1/P4 productive interactions. Five P1 residues (P1.T11, P1.D14, P1.H26, P1.E38, P1.A42) have been shown in a previous mutational study to affect autophosphorylation activity if mutated (32), indicating these are key interfacial residues for productive P1/P4 interactions. As shown in Figure S6 and Table S1, HyRes-predicted *trans-* productive mode (i) and (ii) capture all five P1 residues with >0.1 contact frequency (Figure S6A & S6B and Table S1, labeled in red), except P1.H26 in mode (i). As for P4 residues, P4.S351 (P4.S334), P4.D359 (P4.D342), P4.I388 (P4.I371), P4.R446 (P4.R429) and P4.V485 (P4.V468) in *S. typhimurium* CheA (*E. coli* numberings are shown in the parenthesis) have been shown to interfere P1/P4 autophosphorylation activity when chemically modified (33). Both predicted *trans-* productive modes only capture the P4.D342 site with >0.1 contact frequency while missing the rest (Figure S6A & S6B). Importantly, 24 sites on P1 and P4 have been experimentally determined to be functionally non-perturbing when chemically modified (33) (Table S1 and Figure S7, colored in blue). Both predicted *trans-*productive modes show that these sites are not involved with high contact frequencies except for 3 sites in mode (i) and 2 sites in mode (ii) (see Tabel S1). Taken together, the identified *trans-*productive modes of P1/P4 interactions are largely consistent with available experimental data, accurately distinguishing important P1/P4 interfacial residues from insignificant ones.

Comparing these two predicted productive modes to the previous model (Figure S8), the same P1 helices A and B are involved at the interface, but the P4 binding surface differs from the P1/P4 binding model proposed by Zhang *et al* (31). Of note, P1 interacting interface residues identified from the previous mutational study (32) locate on helix-A and helix-B and are consistent with both Zhang’s productive model and the current modes derived from HyRes simulations. On the other hand, as shown in Figure S8, the α1, α2 and α6 helices of P4 provide the binding surface for P1 in HyRes-predicted productive modes, while the α3, α4 and the ATP-lid of P4 form the binding surface in Zhang’s model. It is possible that multiple *trans-* productive modes, including those identified here and previously, may co-exist to support P1/P4’ phospho-transfer reactions but are functionally affected to different extents by mutations or chemical modifications. More direct experiments such as chemical shift perturbation of P4 may be necessary to further resolve the P1/P4 interaction interface in various functional states.

### Possible non-productive modes of P1/P4 interaction

The P1 COM densities below the P4 domains (“lower” docking densities) observed in HyRes simulations are consistent with the featureless densities captured by previous cryo-ET studies (12, 18). These nonspecific P1/P4 binding configurations do not represent a productive interaction because the P1 domain (and specifically H48) is far from the ATP binding site on P4. As shown in Figure 5, two apparent major non-productive binding modes (labeled (i) and (ii)) can be identified in both ATP-free and ATP-bound CheA systems. This suggests that ATP has little effect on non-productive binding modes of P1 and P4. The non-productive mode (i) has an inter-helix COM distance of ∼22.5 Å and a directional vector angle of ∼140°; the non-productive mode (ii) has an interhelix COM distance of ∼23.5 Å and a directional vector angle of ∼74.5°. Both non-productive modes use the helix-α1 of P4 and the helix-A and helix-D of P1 to form the binding interfaces (Figure 5).

**Figure 5.**
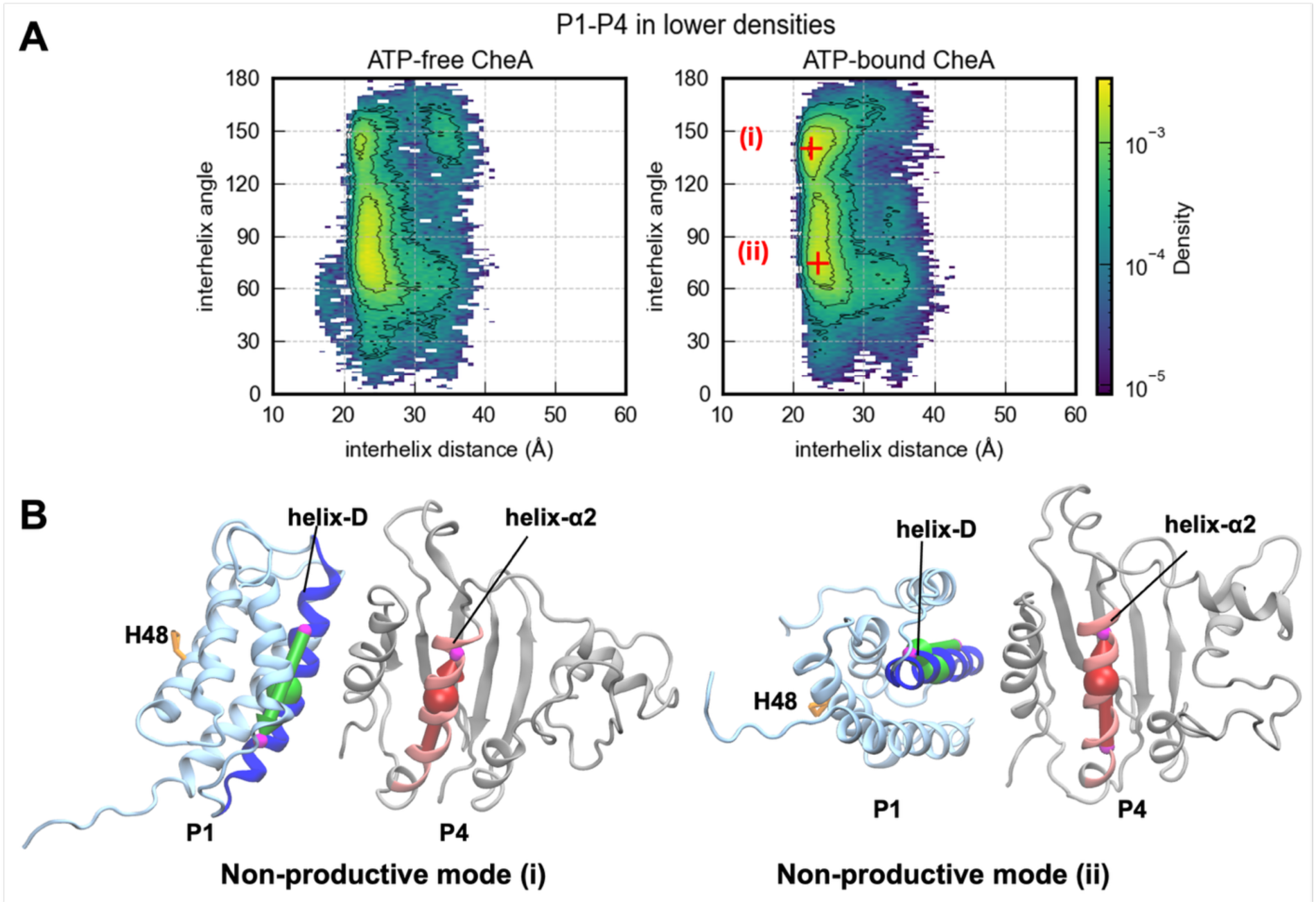
non-productive binding modes from the “lower” docking densities. **A)** Probability distributions of non-productive binding configurations as a function of the COM distance and directional vector angle between helix-D of the P1 domain and helix-α2 of the P4 domain (see panel B). B) Two representative snapshots of the major P1/P4 nonproductive binding modes, corresponding to the cluster center (i) and (ii) (marked by red “+” in panel A) from the ATP-bound CheA state. The directional vectors of the helix-D of P1 and helix-α2 of P4 are shown as green and red arrows, respectively. The COMs of the helix-D and the helix-α2 helix are shown as green and red spheres. H48 is shown as orange sticks.

We further analyzed whether the predicted non-productive P1/P4 interaction modes are consistent with the non-productive binding surfaces proposed from an NMR study of CheA from *T. maritima* (30). Their proposed non-productive binding mode involves *T. maritima* P1 Helix D residues D93, M94 and R97 (corresponding to D96, I97 and E100 in *E. coli*) and *T. maritima* P4 residues E387 and V396 (corresponding to S358 and I367 in *E. coli*), which showed the largest chemical shifts when adding free P1 to a P3P4 construct. We calculated the distances between the COM of D96, I97 and E100 in P1 and S358 and I367 in P4 in both the ATP-bound and ATP-free systems (see Methods). There were no obvious *trans-* or *cis-* contacts between either P4.S358 or P4.I367 and the three Helix-D residues from P1 sampled in both systems among all simulations, meaning non-productive contacts like those previously identified in *T. maritima* were not sampled in the simulation of the *E. coli* CheA dimer (Figure S9). However, it is worth noting that both HyRes-predicted non-productive modes have high contact frequencies for D96, I97 and E100 in helix-D of P1, which aligns well with the NMR study (Figure S10A & S10B). The negatively charged residues mainly interact with a positively charged residue cluster on helix-α1 of P4, namely R335, R338 and R341, instead of S358 and I367 (Figure S10C & S10D). This result indicates electrostatic interactions may dictate how P1/P4 domains form non-productive binding in *E. coli*. These differences in the P4 residues that form the non-productive binding surface for P1 could be due to differences in the sequence of the P4 domain between *E. coli* and *T. maritima* CheA.

Interestingly, the predicted non-productive mode (i) is similar to Cassidy’s non-productive mode (18) (Figure S10C and S10E). The position and orientation of the P1 domain, the P1/P3P4 interacting surface and even positively charged residue cluster (R355, R338 and R341) on the helix-α1 of P4 forming close contacts with P1 are shared in these two proposals. Therefore, HyRes simulations not only recapitulate the previously identified non-productive mode with consistent interfacial interactions but also explore broader domain-domain interaction configurations (Figure 5A) resulting in the alternative non-productive mode (ii).

### Possible auto-inhibitory modes of P1/P1’ interactions

Previous experimental data has suggested that P1/P1’ interactions may reduce autophosphorylation (37, 38), especially if P1/P1’ dimer interactions also involve helix-A and helix-B and shield the H48 phosphorylation site on helix B. Such interactions are found in a parallel P1 dimer (Figure S11B) observed in P1 crystals and consistent with EPR distance measurements (34). It is also observed in an antiparallel P1 dimer (Figure S11C) predicted by AlphaFold-2 (18). Alternatively, P1/P1’ interactions involving the helix-A and helix-D helices may not occlude H48, such as in a previous dimer model involving helices A and D at the dimer interface to account for very slow deuterium uptake in these helices (29). However, the transient nature of the P1 dimers (see below) indicates that the slow deuterium uptake of the A and D helices is likely due to their intrinsic stability as observed in a previous NMR study of the P1 monomer (60). Furthermore, this interface contains many negatively charged residues, making it highly unlikely for Helix-A and Helix-D to form a dimerization interface (Figure S11A). Both the Cassidy (16) and Muok (32) P1/P1’ models also contain several residues of the same charge directly contacting each other (Figure S11B & S11C), which is highly unfavorable.

Here we analyzed the detail of P1/P1’ interactions in both ATP-free and ATP-bound simulation trajectories. About 11-13% of all snapshots sampled have helix-B of P1 and P1’ within a 25 Å COM distance. As shown in Figure S12, P1/P1’ dimer configurations were rarely sampled in the upper docking region in both ATP-bound and ATP-free states, with a similar major minimum at a large helix-B distances of ∼ 60 Å. Instead, possible dimerization occurred primarily in the “lower” docking region (29-36% of the configurations sampled in the “lower” region, see Table S3). In Figure 6, we compare the distribution of P1/P1’ dimers derived from the ATP-free and ATP-bound trajectories. Wide ranges of the interhelix distances (10 - 70 Å) and angles (0 - 180°) were sampled, reflecting a highly dynamic and transient nature of P1/P1’ interaction in CheA dimers. ATP binding to P4 has limited effect on P1/P1’ interactions; both overall distributions and major dimerization modes are highly similar (Figure 6A). Three major P1/P1’ dimerization modes can be resolved, denoted as (i), (ii) and (iii) in Figure 6A right panel. The mode (i) has an interhelix distance and angle of ∼27 Å and 100°, representing a “back-on-face” docking configuration for the two P1 domain (Figure 6B). According to contact frequency maps shown in Figure S13, the interfacial contacts mostly involve the helix-A,B from one P1 and the helix-D,E from the other. The mode (ii) adopts a parallel “face-on-face” configuration with an interhelix distance and angle of ∼15 Å and ∼50°, where the helix-A and helix-B form the dimerization interface (Figure 6B middle panel). The mode (ii) has the closest interhelix distance and is semi-symmetrical with a parallel packing style reminiscent of the Muok *et al’*s model (34) (see Figure S11B). The calculated contact map is almost symmetrical (Figure S12) and shows oppositely charged pair interactions involving R45 and E18/E15. Largely, the mode (ii) is similar to Muok *et al*’s model, which has an interhelix (helix-B) distances and angles of 11.8 Å and 21°, in that both use helix-A,B as the dimerization interface and have the substrate site H48 buried on the dimerization interface (Figure 6B & Figure S14B). The mode (iii), while sharing roughly the same interhelix angle (∼50°) with the mode (ii), has a slightly larger interhelix distance of ∼23 Å. The contact map of the mode (iii) is similar to that of the mode (ii) but with smaller contacting frequencies (Figure S13).

**Figure 6.**
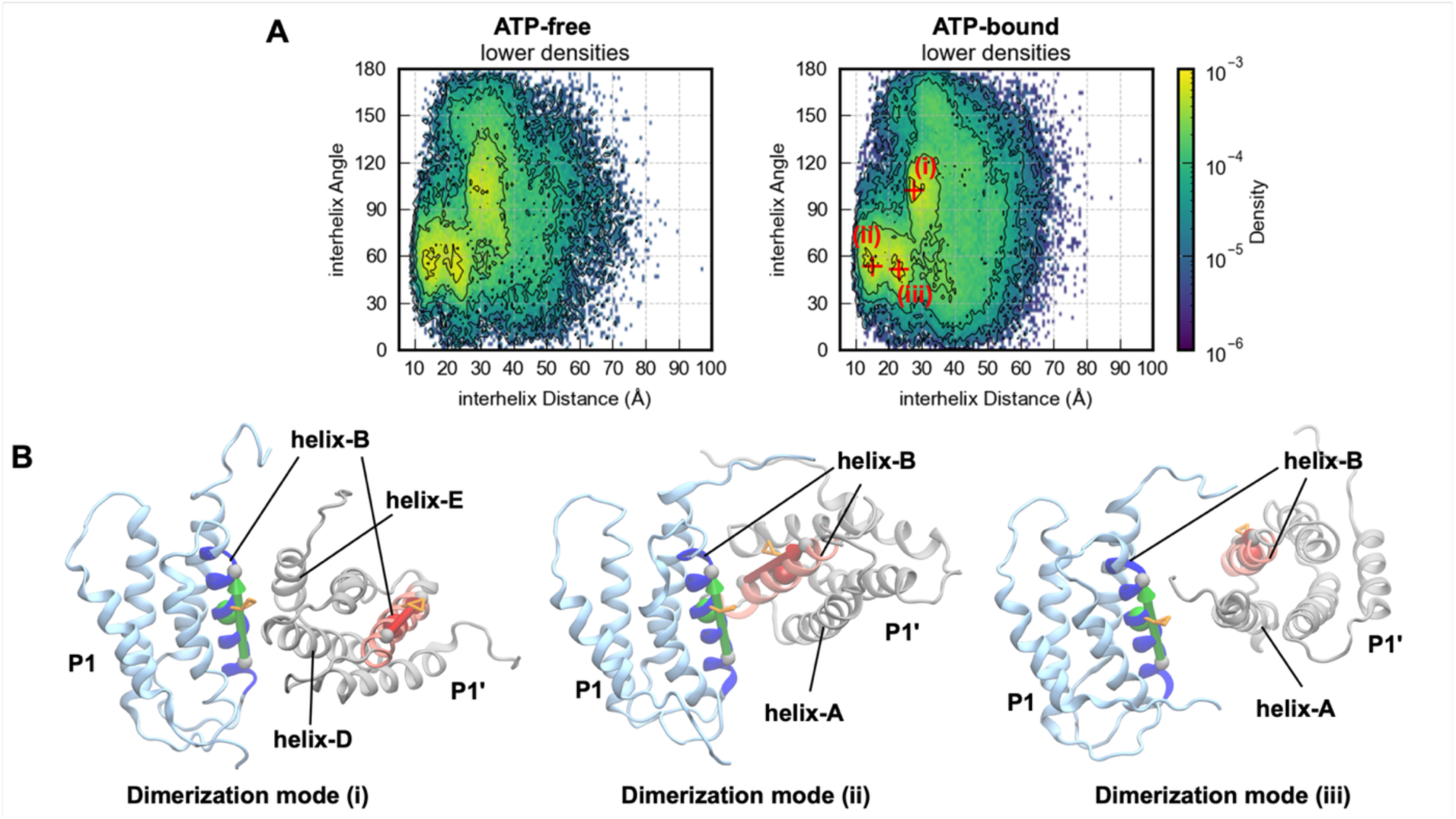
P1/P1’ domain dimerization in the CheA dimer. **A)** The density contour plot of all snapshots from ATP-free and ATP-bound CheA simulations as a function the COM distance and directional vector angle between helix-Bs of the P1 and P1’ domains (see Methods). The centers of major binding modes (i)-(iii) are marked using “+” on the right panel. **B)** Representative snapshots of the P1 dimerization modes. The directional vectors of two helix-B from P1/P1’ are shown with green and red arrows, with the two helix-Bs colored in blue and red. H48 is shown as orange sticks. The snapshots are aligned using the P1 domain with interfacial helices annotated.

To further study P1 dimerization modes in the absence of other domains (P2-P5), we performed 2 µs HyRes simulations of two isolated P1 domains (P1/P1’) to derive the distribution of angles and COM distances between the helix-B from the two P1 domains (see Methods). The results reveal the two most major dimerization modes, denoted as mode (1) and (2) (Figure S14). Interestingly, modes (1) and (2) share similar interhelix distances with the mode (i) and (ii) respectively, but with the interhelix angle increased by ∼30°. The difference in preferred interhelix angles is likely due to the crowded and restrained environment in CheA dimers as well as competition from P1/P4 interactions. Therefore, P1 dimerization modes from the whole CheA systems and from the two P1 domain simulations are largely consistent and support the experimental observations that P1 domains in the CheA dimer can sequester each other effectively in the “lower” docking region for auto-inhibition.

## Discussion

Modulation of autophosphorylation activity of the histidine autokinase CheA is a critical step during chemotaxis signaling in motile bacteria. Due to the inherent mobility of the P1 and P2 domains in CheA, structural studies so far have struggled to capture their docking modes on other domains (12, 16, 18, 28). Other experimental techniques, including NMR (30), chemical labeling (33) and mutational screening (32), have yielded important but limited evidence for the multiple possible docking interactions between CheA domains. Herein, we adopt a bottom-up approach, that is, extensive coarse-grained simulations of the CheA dimer with or without ATP bound in P4, to systematically investigate all possible interactions between domains P1 and P2 and the kinase core (P3P4P5). Preferential docking regions of P1 on P4 were identified and found to be largely similar for both the ATP-bound and ATP-absent systems (Figure 2). The P1 domain seems to interact with 2 main regions of P4: the “upper” region that includes the productive interface, and the “lower” region that we found did not contain the previously proposed non-productive interface. The supplemental movies illustrate the transient nature of these interactions, in which ∼60% of the frames have at least one P1 in the upper region, and ∼30-50% of the frames have at least one P1 in the lower region.

The “upper” region of P1/P4 interactions changes when ATP is bound to P4: the *cis-* upper density disappears, and the trans upper density changes to include productive binding modes. Past studies have established that CheA is phosphorylated in *trans-* (P1/P4’ or P1’/P4) interactions, but have not previously noted a role for ATP in directing the trans interaction. Crane and coworkers proposed, based on a crystal structure of P3-P4 that includes part of the P2-P3 linker and bound AMPPCP, that interactions between positively charged residues on the P2-P3 linker and negatively charged residues in the P4 domain could be important for directing P1 towards a trans-interaction with P4’. We suggest that ATP binding changes the P4 conformation to expose those negatively charged P4 residues that interact with the P2-P3 linker to direct P1 towards trans P1/P4 interactions. Our study also shows that productive P1/P4 interactions do not occur in the absence of ATP. This is consistent with kinetic studies that indicate an ordered mechanism, with ATP binding before P1, and biolayer interferometry that demonstrated P1 does not bind P3P4P5 until an ATP analog (AMPPCP) is added (61). In addition, HDX-MS data shows that addition of AMPPCP leads to slower deuterium uptake in regions around the P1 substrate site and the P4 catalytic core, consistent with AMPPCP promoting the productive P1/P4’ interaction (29). There have been two proposals for the role of ATP in promoting P1 binding. The ATP lid of P4 was previously proposed to change from disordered to helical upon nucleotide binding (36), which was proposed to create the binding surface for productive interaction with P1 (61). However, the ATP lid exhibits rapid deuterium uptake even in the presence of AMPPCP (29). Furthermore, since the ATP lid in all our simulations was modeled as a loop, a folded ATP lid is not required to see the ATP effect in promoting the P1/P4’ productive interaction. MD simulations instead proposed that ATP binding causes the ATP lid to shift position to an open conformation that allows P1 to bind, and yielded a model for the P1/P4 productive interaction (Zhang model) in which the P1 binding surface involves primarily helices α3 and α4, with only two residues from the ATP lid (31). The two productive binding modes found in our simulations involve a different surface of P4 than the Zhang model. Both HyRes productive binding modes involve the α1, α2, and α6 helices of P4, and this difference could change the expected mechanism of regulation of CheA. For example, signal propagation may come through the P3-P4 hinge into the connected α1 helix of P4 to cause slight changes that favor the productive interaction. Thus our simulation study has identified multiple roles for ATP in favoring the productive P1/P4 interaction and yielded two additional possible models for that interaction.

Although we did not observe the previously proposed non-productive interaction between Helix D of P1 and *E. coli.* residues S358 and I367 of P4 in our simulations, the “lower” COM density for P1 likely represents conformations that similarly sequester and inhibit productive interactions, since the substrate site of P1 in those regions is similarly inaccessible to the ATP-binding pocket in P4. Although the most likely P1/P1’ dimer interactions would also be inhibitory, as these bury the phosphorylation site at the dimer interface, this is not likely to regulate CheA activity, since possible dimers are found primarily in the lower COM density that is already a nonproductive interaction. P1/P1’ interactions are found to be rare relative to P1/P4 interactions, so the previously observed modest contribution of P1 to CheA dimer affinity (62) is likely due to the more frequent P1/P4 interactions. Our simulations are consistent with previous evidence (28) that the amorphous density below P3 in cryo-ET density maps of the CSU complex (12, 18) contains P1 and P2, and further suggest that only ∼10% of the P1 is dimeric.

Since the role of P2 in CheA is simply to bind to response regulator proteins such as CheY, there are no necessary interactions between the P2 domain and other domains of CheA during the catalytic cycle of autophosphorylation or phosphoryl transfer. In a previous solution NMR study of full-length CheA (63), only the P2 domain was visible in the spectra, suggesting that its flanking linkers enable it to move much more rapidly than the rest of CheA (resonances of the 142 kDa CheA dimer are broadened by its slow rotation). Therefore, P2 has very little interaction with other domains. Although the P1 domain is connected to CheA by the same flexible linkers, it was not visible in the NMR spectra, due to significant interactions that restrict its mobility. This picture is consistent with the effect of ATP in our simulations: the P2 distribution is unaffected since it is not interacting with P4, but the P1 distribution is affected, consistent with its significant interactions with P4.

Our study provides an in-depth bottom-up investigation of the dynamic domain-domain interactions in the CheA autokinase dimer. The binding modes between P1/P4 domains derived from our simulations lay the groundwork for further investigation of autophosphorylation regulation as well as downstream interactions with CheY. Future studies of these domain-domain interactions in the context of the CSU complex (including the receptor cytosolic domain and CheW) using coarse-grained simulations may provide a more complete picture of the domain dynamics and interactions of CheA that regulate its autophosphorylation.

## Acknowledgements

We thank Sandy Parkinson for coordinates for the Zhang productive model. This work is supported by NIH through grants R35 GM144045 (to Chen) and R01-GM120195 (to Thompson). JH was partially supported by a fellowship from the University of Massachusetts as part of the Chemistry-Biology Interface Training Program (National Research Service Award T32 GM139789). KAW was partially supported by National Research Service Award T32 GM008515 from the National Institutes of Health.

## Author contributions

Thompson and Chen conceptualized the idea. Huang conducted computational modeling. simulation, and analysis. Huang, Wahlbeck, Thompson and Chen interpreted data, drafted and revised the manuscript.

## Additional information

**Extended data** is available for this paper at https://doi.org/xxxx

## Supplementary information

The online version contains supplementary material available at https://doi.org/xxxx

