## Supplemental tables and figures for "Dynamic domain interactions encode possible CheA autophosphorylation mechanisms revealed by coarse-grained simulations"

### Supplementary Movie Captions

**Movie S1** One example trajectory (2 μs) from the ATP-free CheA system. The CheA dimer is shown as cartoon representations with P1-P5 colored blue, gray, orange, pink and olive. Two long flexible linkers (connecting P1/P2 and P2/P3) are colored green. The restraining scheme applied to the system is thoroughly described in Methods. Briefly, P1, P2 and P4 domains were internally restrained to keep their 3D structures intact while P3 and P5 were positionally restrained with respect to the original model.

**Movie S2** One example trajectory (2 μs) from the ATP-bound CheA system. The CheA dimer is represented the same as in Movie 1, while the HyRes-resolution ATP is shown as ball-and-stick with beads forming the adenine group, the ribose ring, and phosphate groups colored in blue, cyan, and red, respectively. The same restraining scheme as in the ATP-free system was adopted, and we added additional distance restraints between ATP and P4 residues (see Methods for details).

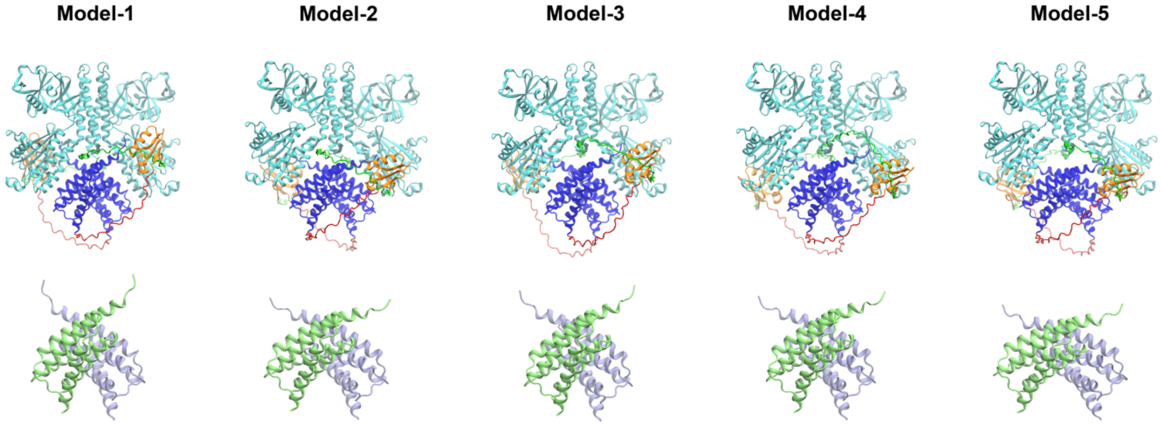


**Figure S1. Five AlphaFold2 models of the CheA dimer.** The models were predicted by mmseqs2 and then refined by Rosetta (see Methods)**. A**ll models are aligned to C$\alpha$ atoms of the P3, P4 and P5 domains of Model-4, which ranks the highest based on the AlphaFold2 prediction. P3-5 C$\alpha$ RMSD to Model-4 is 1.6 Å, 1.1 Å, 0.7 Å, 1.2 Å for Model-1, Model-2, Model-3 and Model-5, respectively. Although there are some positional shifts of P1 and P2 among the models, the internal structures of these domains are highly similar to available PDB structures (C$\alpha$ RMSDs for all 5 models to pdb:2LP4 is 0.37 - 0.48 Å for P1 and 1.39 – 1.45 Å for P2). The P3, P4 and P5 domains are colored in cyan, and the P1 and P2 domains are colored in blue and orange respectively in the upper panel. The disordered linkers connecting P1/P2 and P2/P3 are colored red and green. The P1/P1' domains from each model are separately shown in the lower panel and are colored in lime and ice blue.

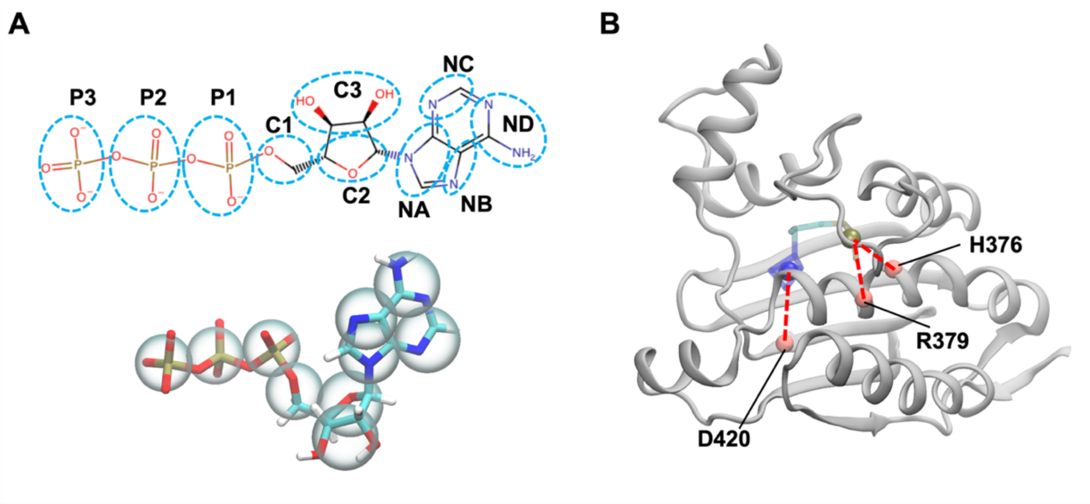


**Figure S2. Modeling bound ATP. A)** The mapping scheme of the HyRes ATP molecule. **B)** The distance restraints set up on the ATP molecule in ATP-bound CheA system. Only the P4 domain of the CheA is shown as cartoon. The HyRes ATP is shown using sticks. The blue and tan spheres are the COMs of the ATP headgroup and three tail phosphate groups respectively. For harmonic distance restraints (red dashed lines), a $K_{constant}$ = 400 KJ/mol/nm$^2$ was used and the restraining distances between ATP COMs and C-α atoms of D420, R379 and H376 are 10, 8, 8 Å respectively.

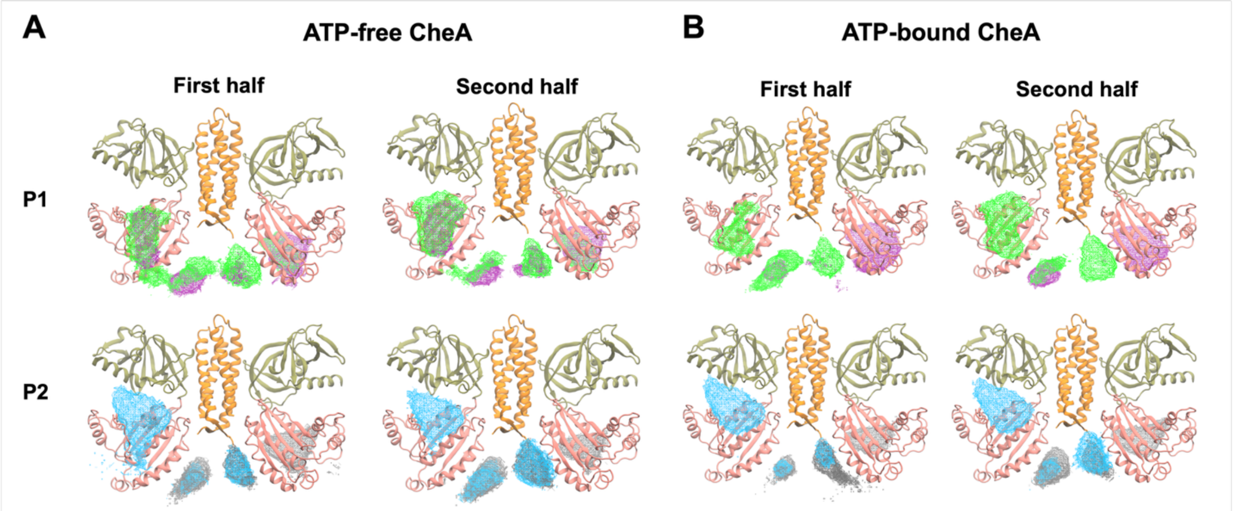


**Figure S3. Convergence of COM densities in the ATP-free CheA (A) and ATP-bound CheA (B) system.** All six trajectories in each system were split into the first halves and second halves. Then, the P1 and P2 COM densities were calculated for all combined first halves and second halves respectively. All density maps are shown using an iso-value of 6 e/Å$^2$.

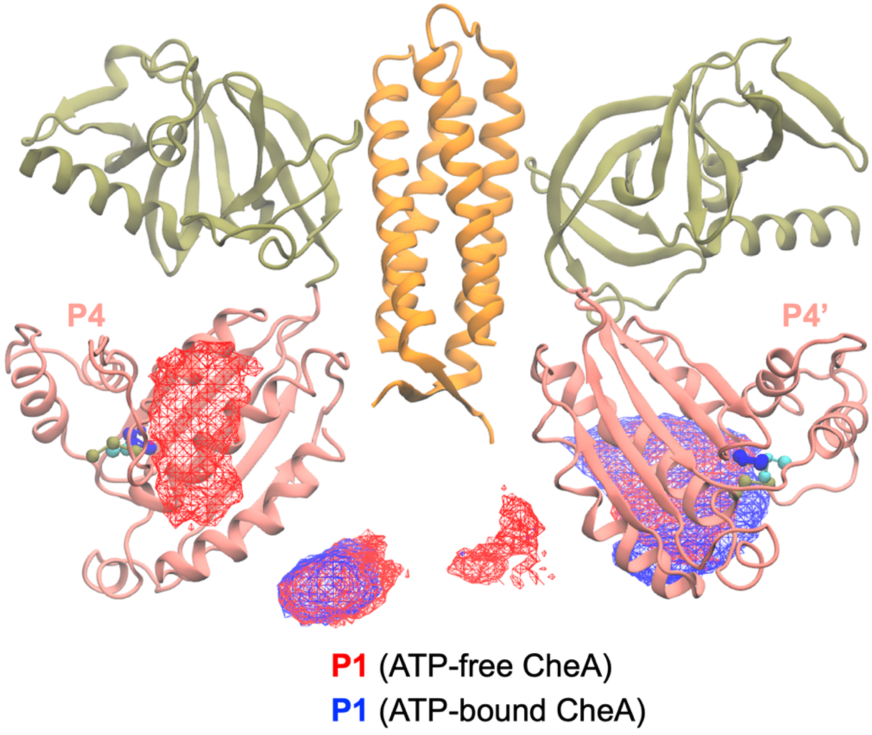


**Figure S4. Overlay of the P1 densities from the ATP-free (red) and ATP-bound (blue) systems.** Only the P1 (not P1') densities of both systems are shown at an iso-values of 12 e/Å$^2$.

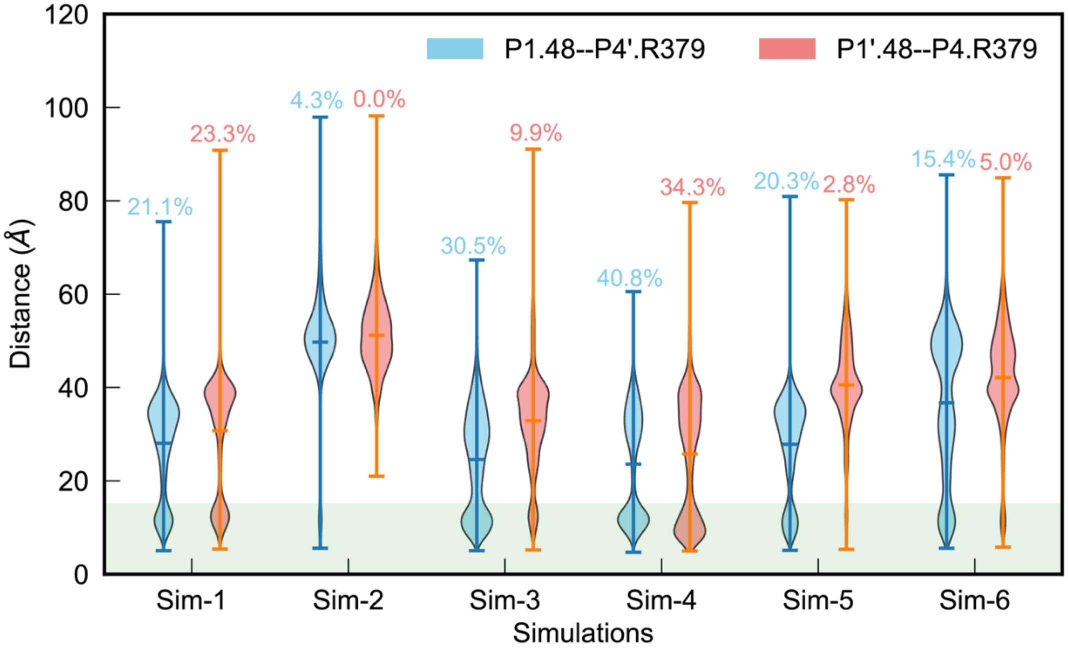


**Figure S5. Productive-like contacts in the ATP-bound state.** The distances between P1.H48 and P4'.R379 (cyan) and between P1'.H48 and P4.R379 (salmon) are calculated and plotted separately as violin plots. A cutoff distance of 15 Å (green shaded region) was used to determine the probability of forming possible *trans*- productive contacts, which are marked above each violin plot.

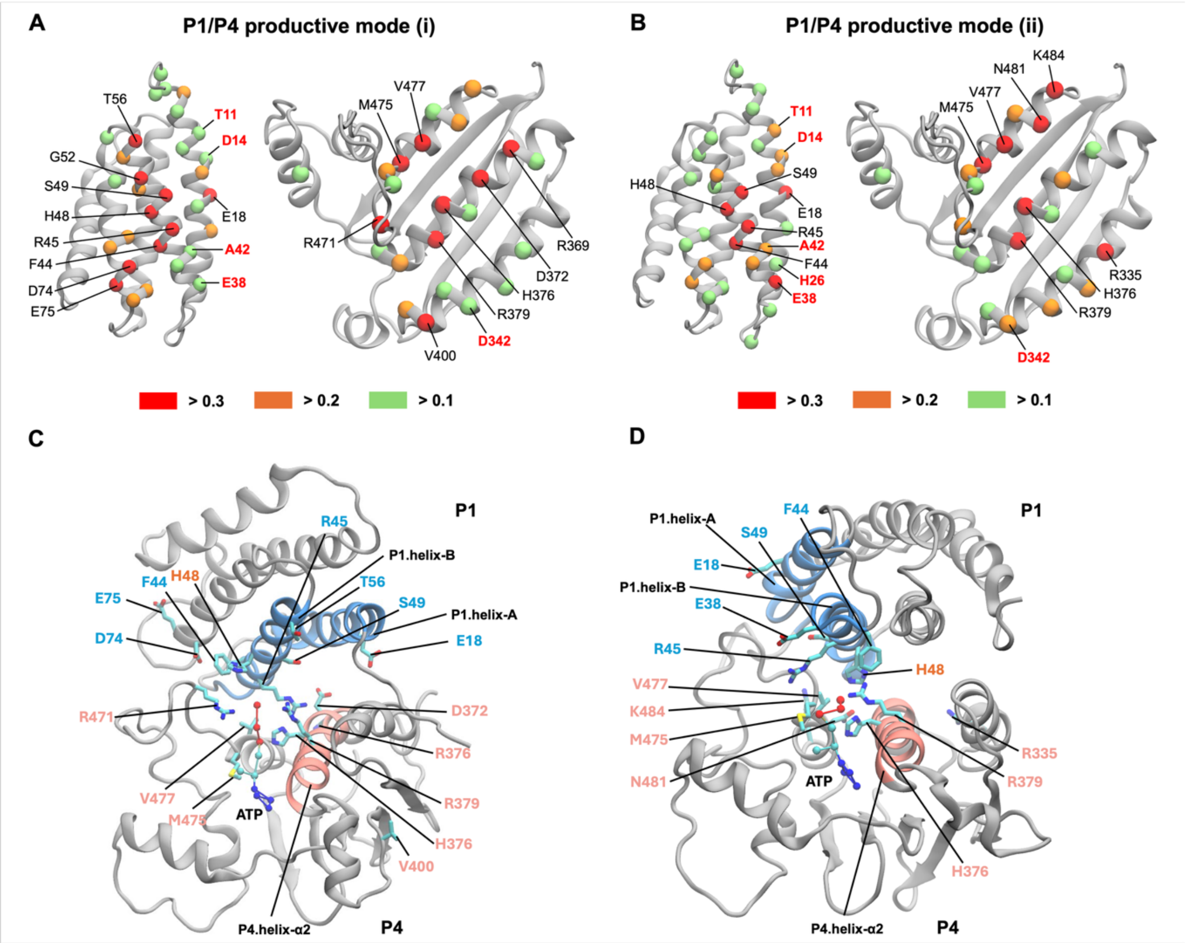


**Figure S6. Contact frequency analysis of the two identified *trans-* P1/P4 productive binding modes. A)** and **B)** P1 and P4 residues with high contact frequencies from the productive mode (i) and mode (ii). C-α of residues with contact frequencies larger than 0.3 are shown as red spheres and labeled in black; larger than 0.2 and less than 0.3 are shown as orange spheres; larger than 0.1 and less than 0.2 are shown as green spheres. Contact residues identified by experimental studies to be important to the productive interaction are labeled in red. **C)** and **D)** Residues at the P1/P4 contacting interface in the productive mode (i) and (ii), respectively. Interfacial residues are shown if they are contact frequencies are larger than 0.3 (red sphere residues in A) and B)). P1 and P4 domains are shown as gray cartoon representations, except P1 helix-A and helix-B (colored blue) and P4 helix-α2 (colored salmon). Residues at the contact interface are shown as sticks with C, O and N atoms colored using cyan, red, and blue respectively. Residues on P1 are labeled in light blue and residues on P4 are labeled in pink. The ATP molecule is shown as balls and sticks in the Hyres resolution. The substrate site H48 is labeled orange.

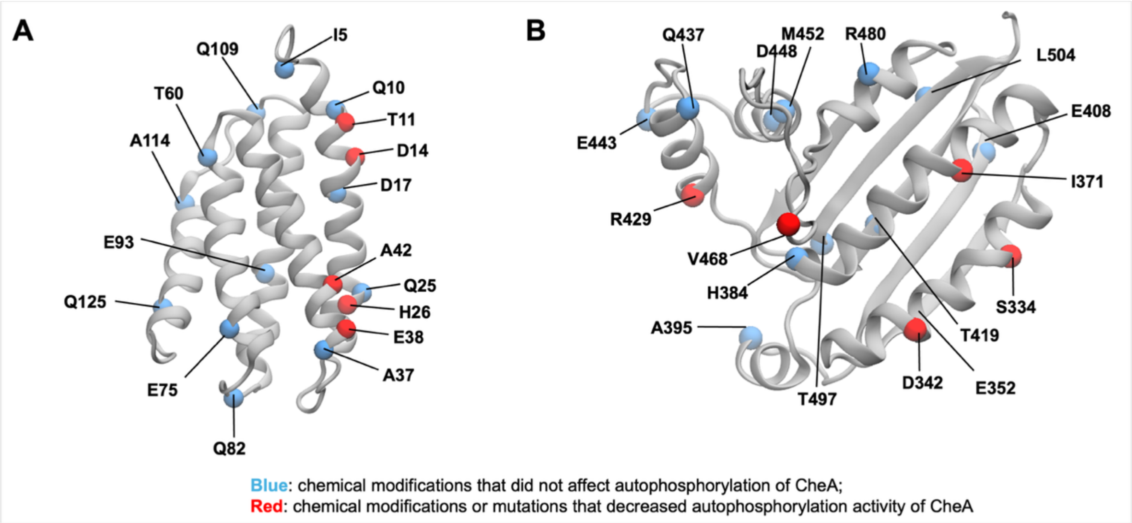


**Figure S7. Experimentally verified mutation or chemical modification sites in P1 (A) and P4 (B)**. Sites that affect or do not affect autophosphorylation activities of CheA are colored in red and blue, respectively.

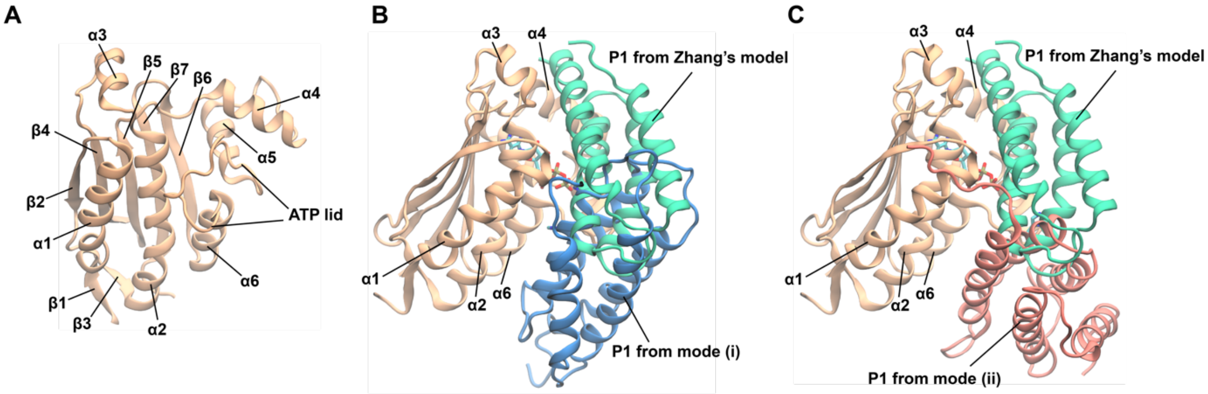


**Figure S8. Comparison of HyRes-predicted P1-P4 *trans-* productive modes with the model from Zhang et al (1). A)** Secondary elements of the P4 domain, extracted from Model-4 of AlphaFold2 predictions (Figure S1). **B-C)** Comparison of HyRes-predicted P1-P4 docking modes (i) and (ii) with the one from Zhang et al (1). The P4 domains were used for structural alignment, and the view was rotated 90° horizontally from the view in A) for clarity.

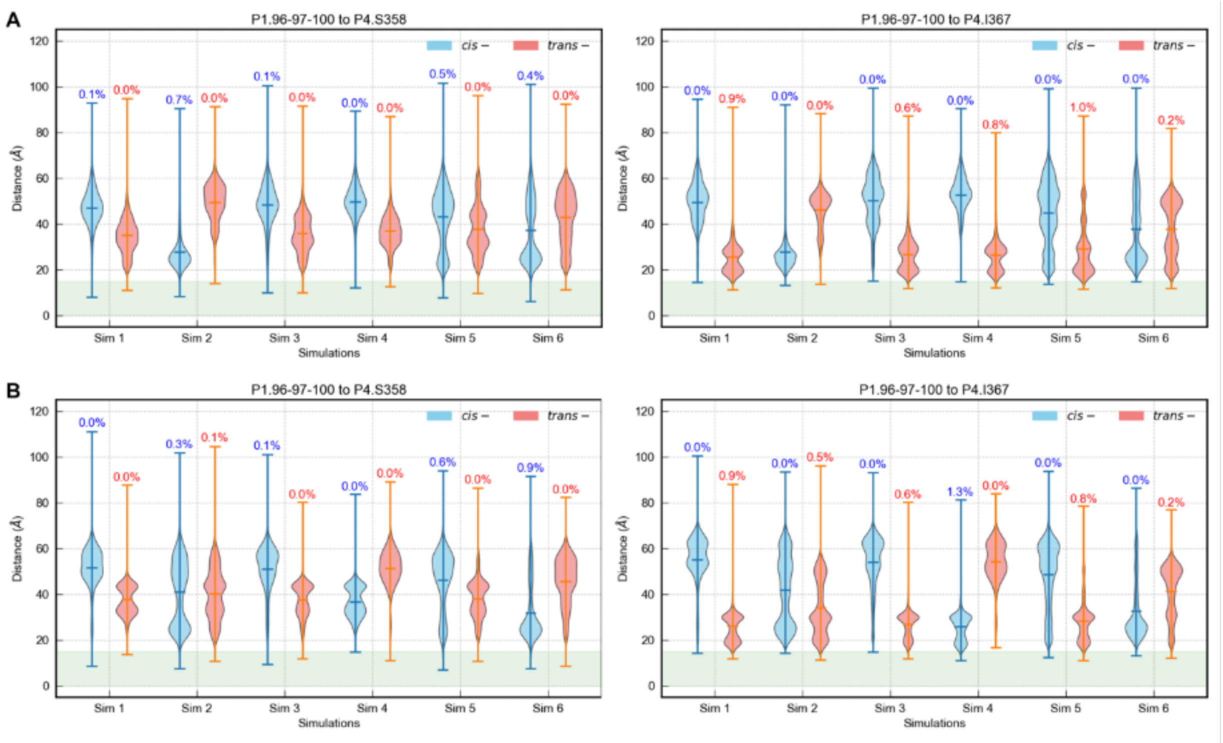


**Figure S9. Nonproductive contact mode analysis for the ATP-bound (A) and ATP-absent (B) system simulations.** The distance distributions between P4.S358 or P4.I367 and COM of P1.96, 97 and 100 residues were calculated from the HyRes simulations (See Methods for calculation details) for both *trans*- and *cis*- distances in the CheA dimer and colored salmon and cyan respectively. A cutoff of 15 Å was used to calculate the fractions of possible contacts, which were reported as on top of each violin distribution.

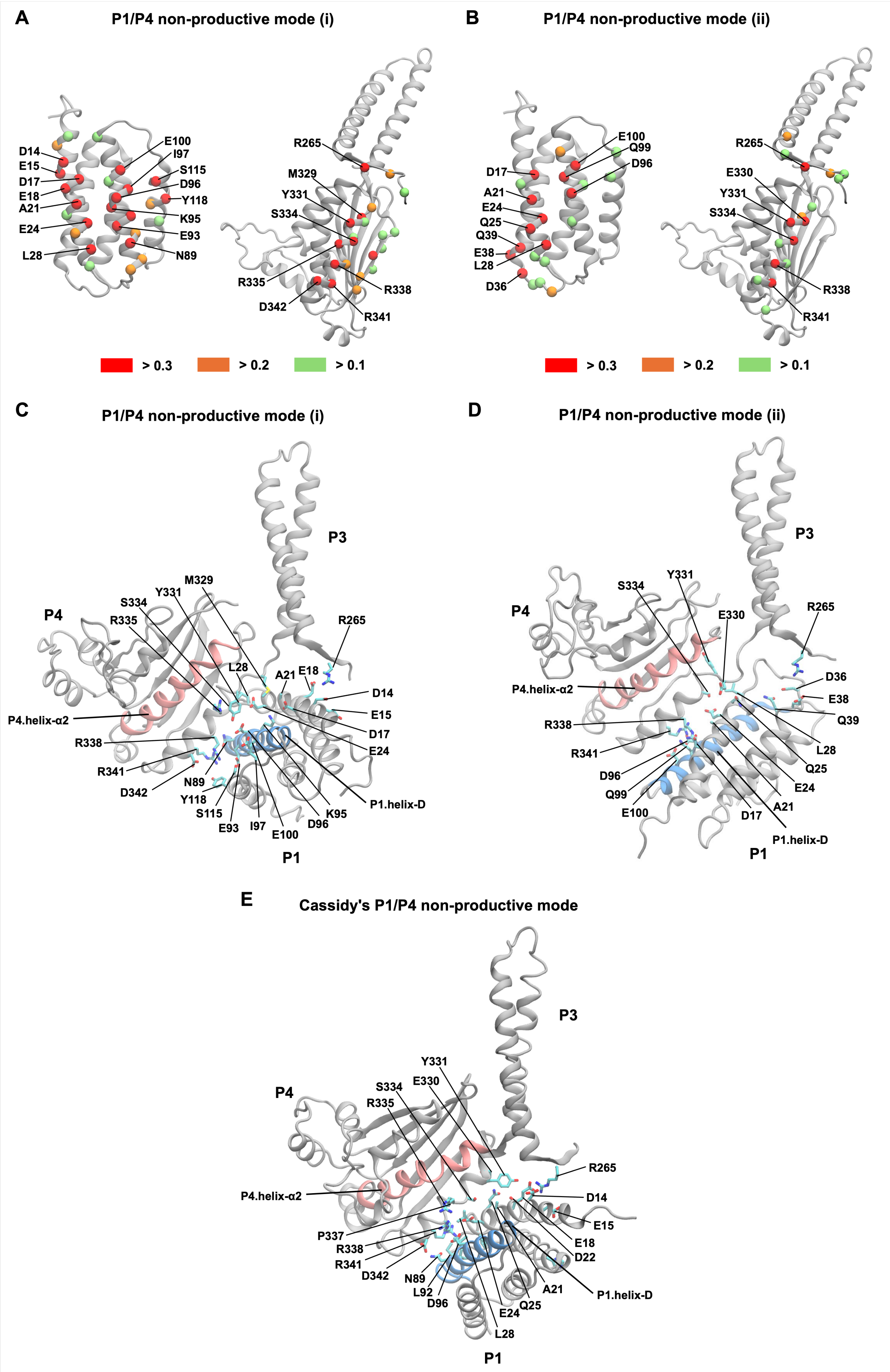
A
P1/P4 non-productive mode (i)
D14
E15
D17
E18
A21
E24
L28
E100
I97
S115
D96
Y118
K95
E93
N89
R265
M329
Y331
S334
R335
D342
R338
R341
> 0.3
> 0.2
> 0.1
B
P1/P4 non-productive mode (ii)
D17
A21
E24
Q25
Q39
E38
L28
D36
E100
Q99
D96
R265
E330
Y331
S334
R338
R341
> 0.3
> 0.2
> 0.1
C
P1/P4 non-productive mode (i)
P3
P4
M329
Y331
S334
R335
R265
L28
A21
E18
D14
E15
D17
E24
P4.helix-α2
R338
R341
D342
N89
Y118
S115
E93
I97
E100
D96
K95
P1.helix-D
P1
D
P1/P4 non-productive mode (ii)
P3
P4
S334
Y331
E330
R265
D36
E38
Q39
P4.helix-α2
R338
R341
D96
Q99
E100
L28
Q25
E24
A21
D17
P1.helix-D
P1
E
Cassidy's P1/P4 non-productive mode
P3
P4
Y331
S334
E330
R335
R265
D14
E15
E18
D22
P4.helix-α2
P337
R338
R341
D342
N89
L92
D96
A21
Q25
E24
L28
P1.helix-D
P1

**Figure S10. Contact frequency analysis of the two identified P1/P4 non-productive binding modes and Cassidy's non-productive mode. A)** and **B)** P1, P3, P4 residues with high contact frequencies from the non-productive mode (i) and mode (ii). C-α of residues with contact frequencies larger than 0.3 are shown as red spheres; larger than 0.2 and less than 0.3 are shown as orange spheres; larger than 0.1 and less than 0.2 are shown as green spheres. **C) and D)** Residues on the contacting interface of P1/P3P4 in the non-productive mode (i) and (ii) identified from the ATP-bound system simulations. Interfacial residues are shown if they are contact frequencies are larger than 0.3 (red sphere residues in A) and B)). P1 and P4 domains are shown as gray cartoon representations, except the helix-D of P1 (colored blue) and the helix-α2 of P4 (colored salmon). Residues on the contact interface are shown as sticks with C, O and N atoms colored using cyan, red, and blue respectively. The ATP molecule is shown as balls and sticks in the Hyres resolution. The substrate site H48 is labeled orange. **E)** Residues on the contacting interface of P1/P3P4 in the non-productive mode from Cassidy's (2). Coloring and labeling schemes are the same with **B)** and **C)**. Residues that are within 4.5 Å to other domains in the interface are shown.

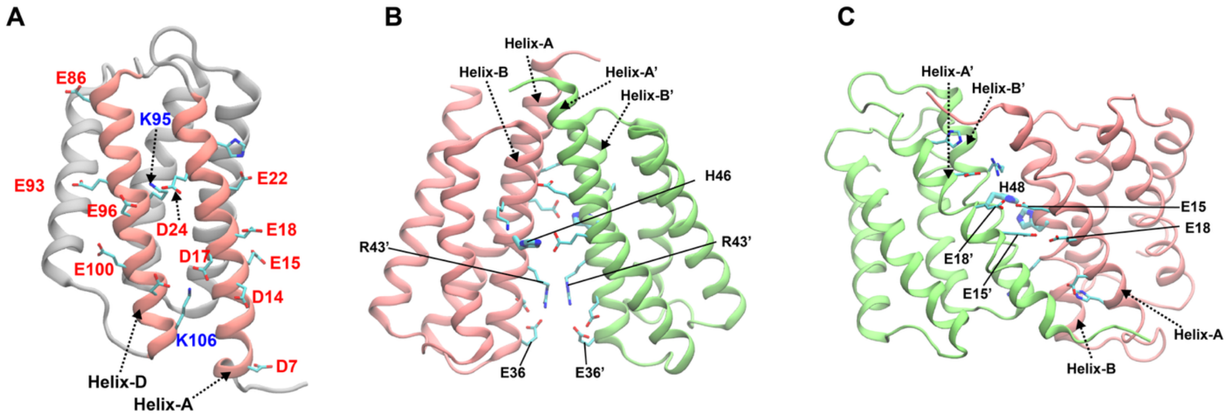


**Figure S11. Charged residue distribution analysis of existing P1/P1' models. A)** Charged residues on the helix-A and helix-D of the P1 domain. *E. coli* residue numbering is used. **B)** The parallel non-productive P1/P1' dimer model from Muok et al (3). Electrostatically frustrated charged residues are labeled. *T. maritima* numbering is used. **C)** The antiparallel non-productive P1/P1' dimer model from Cassidy et al (2). Electrostatically frustrated charges are labeled. *E. coli* residue numbering is used.

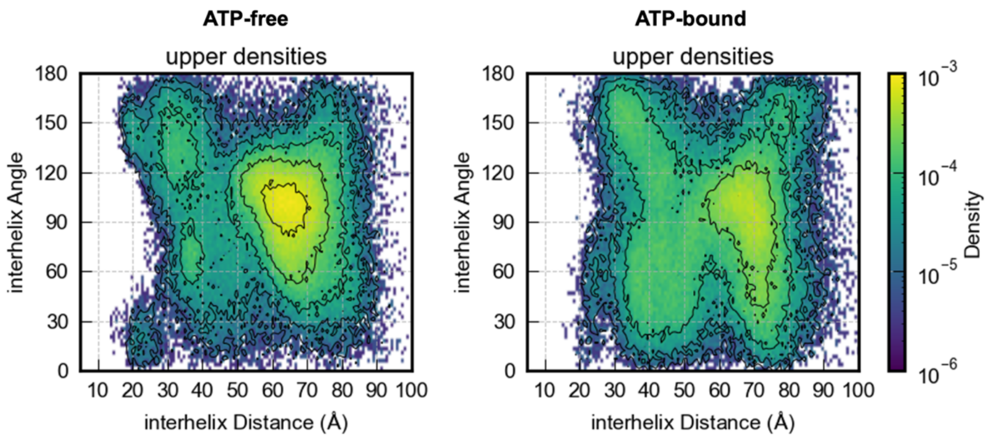


**Figure S12. P1/P1' dimer is rarely sampled in the "upper" docking regions in ATP-free and ATP-bound CheA dimer system simulations.** All snapshots with P1 located in the "upper" docking from all simulations (6*8 μs) were characterized in the left panel using the distance and angle between two helix-B of P1/P1' domains (see Methods).

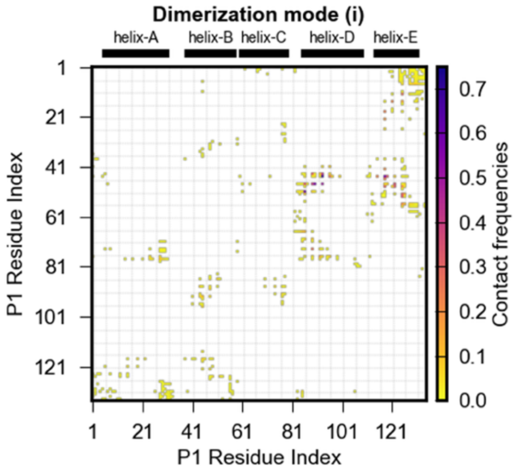


| P1/P1’ Res Index | Contact frequency |
|---|---|
| R45/E93 | 0.70 |
| K51/D86 | 0.62 |
| R45/Y118 | 0.59 |
| H48/L90 | 0.58 |
| H48/E93 | 0.50 |
| F44/E93 | 0.41 |

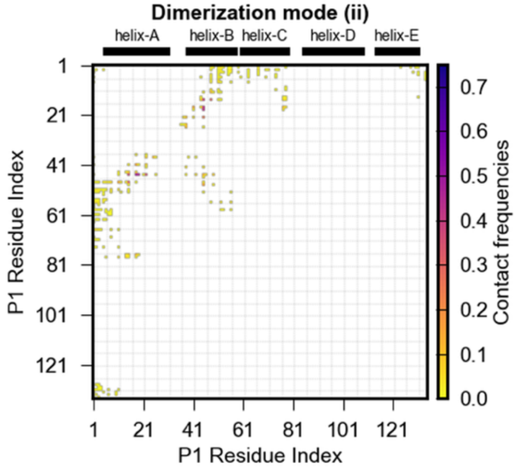


| P1/P1’ Res Index | Contact frequency |
|---|---|
| R45/E18 | 0.50 |
| R45/L19 | 0.46 |
| L19/R45 | 0.42 |
| E18/R45 | 0.41 |
| E15/R45 | 0.39 |
| R45/E15 | 0.38 |

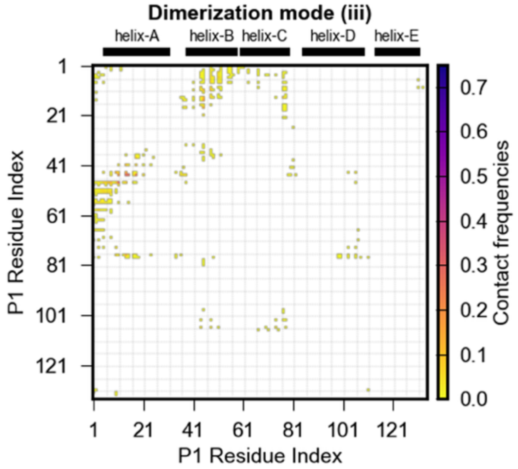


| P1/P1’ Res Index | Contact frequency |
|---|---|
| R45/D14 | 0.29 |
| H48/T11 | 0.28 |
| R45/E15 | 0.27 |
| R45/E18 | 0.25 |
| R45/T11 | 0.18 |
| E15/R45 | 0.16 |

**Figure S13 Contact frequency map of the major P1/P1’ dimerization modes identified from ATP-bound CheA simulations.** Tables on the right show the top highest contact frequencies from those modes.

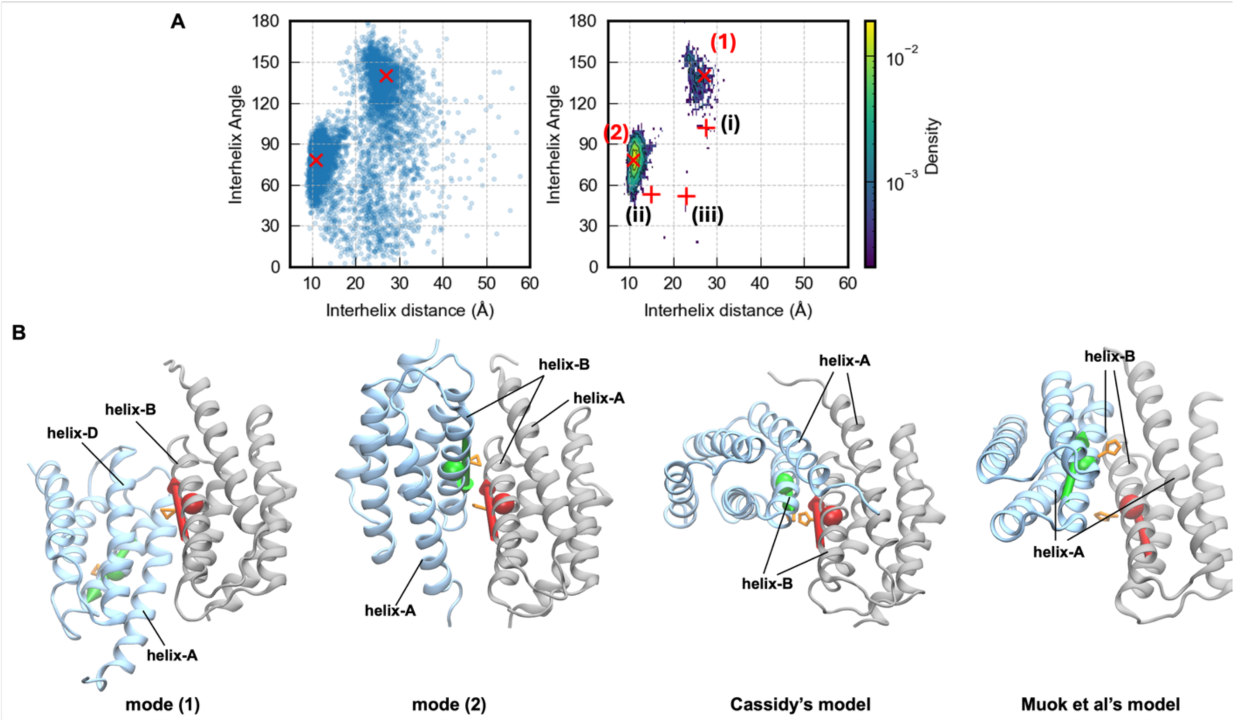


**Figure S14 P1 dimerization modes from the HyRes simulation of two P1 domains. A)** Distribution and density plot of all sampled snapshots from the simulation characterized by interhelix (helix-B) COM distances and directive vector angles between two P1 domains. The major modes (mode (1) and mode (2)) are marked as “x”, while the three modes (i)-(iii) identified from the whole CheA simulations are marked as “+” and labeled for comparison. **B)** Representative snapshots for the mode (1) and (2) labeled the panel A, along with P1 dimer models from Cassidy’s (2) and Muok et al’s (3). Directive vectors of helix-B for P1 domains are drawn as green and red vectors (arrows), while COM of helix-B are shown as green and red spheres. Substrate site H48 (*E. coli*) or H46 (*T. maritima*) are shown as orange sticks. The major (64,2%) binding mode (1) has a helix-B distance of ~10.8 Å and a helix-B angle of ~79.8°, while the minor (12.6%) binding mode (2) has a helix-B distance of ~ 27 Å and a helix-B angle of ~ 140° (Figure 6B). Both of these dimerization modes are antiparallel. The P1/P1’ domains in the major binding mode (1) use helix-A and helix-B to form the dimerization interface, which places the substrate site H48 at the dimer interface and inaccessible for phosphorylation. In contrast, the minor binding mode (2) packs helix-A and helix-D of one P1 domain against helix-B' and helix-C' of the other P1’ domain.

**Table S1.** Residue positions that have been identified to be functionally perturbing (red) or non-perturbing (blue) based on mutational analysis (4) or chemical modifications of the P1 domain (5) (also see Figure S6). Check marks indicate residues found to have >0.1 probabilities of engaging in P1/P4 interactions in productive modes (i) and (ii) (also see Figure S7).

| Sites | Mode (i) | Mode (ii) |
|---|---|---|
| P1.T11 | ✓ | ✓ |
| P1.D14 | ✓ | ✓ |
| P1.H26 | | ✓ |
| P1.E38 | ✓ | ✓ |
| P1.A42 | ✓ | ✓ |
| P4.S334 | | |
| P4.D342 | ✓ | ✓ |
| P4.I371 | | |
| P4.R429 | | |
| P4.V468 | | |
| P1.I5 | ✓ | |
| P1.Q10 | | |
| P1.D17 | | |
| P1.Q25 | | |
| P1.A37 | | |
| P1.S60 | | |
| P1.E75 | | ✓ |
| P1.Q82 | | |
| P1.E93 | | |
| P1.Q109 | | |
| P1.A114 | | |
| P1.Q125 | | |
| P4.E352 | | |
| P4.H384 | ✓ | |
| P4.A395 | | |
| P4.E408 | | |
| P4.T419 | | |
| P4.Q437 | | |
| P4.E443 | | |
| P4.D448 | | |
| P4.M452 | | |
| P4.R480 | ✓ | ✓ |
| P4.T497 | | |
| P4.L504 | | |

**Table S2 Sequence alignment to map P1-P4 docking sites from other species to E. coli numbering.** The UNIPROT accession records (P07363, Q56310, P09384) of the CheA from *Escherichia coli, Thermotoga maritima, Salmonella Typhimurium* are used to perform multisequence alignment using the T-coffee webserver (https://tcoffee.crg.eu/). P1-P4 interaction sites identified from previous studies: Nishiyama et al (4) and Zhang et al (1), Hamel et al (6), and Miller et al (5), are highlighted with green, yellow and magenta colors respectively.

```
sp|P07363|CHEA_  MSMDISDFYQTFFDEADELLADMEQHLLVLQPEAPDAEQLNAIFRAAHSIKGGAGTFGFSVLQ
sp|Q56310|CHEA_  MM---EEYLGVFVDETKEYLQNLNDTLLELEKNPEDMELINEAFRALHTLKGMAGTMGFSSMA
sp|P09384|CHEA_  MSMDISDFYQTFFDEADELLADMEQHLLDLVPESPDAEQLNAIFRAAHSIKGGAGTFGFTILQ

cons             *    .::  .*.**:.* * :::: ** *  :. * * :*  *** *::** ***:**: :


sp|P07363|CHEA_  ETTHLMENLLDEARRGEMQLNTDIINLFLETKDIMQEQLDAYKQSQEPD-AASFDYICQALRQ
sp|Q56310|CHEA_  KLCHTLENILDKARNSEIKITSDLLDKIFAGVDMITRMVDKIVSEGSDDIGENIDVFSDTIKS
sp|P09384|CHEA_  ETTHLMENLLDEARRGEMQLNTDIINLFLETKDIMQEQLDAYKNSEEPD-AASFEYICNALRQ

cons             :  * :**:**:**..*:::.:*::: ::   *:: . :*   .. . * . .:: :.::::.


sp|P07363|CHEA_  LALEAKGETPSAVTRLSVVAK--SEPQ---DEQSRSQSPR----------------------
sp|Q56310|CHEA_  FASSGK-EKPSEIKNETETKG--EEEH-----KGESTSNEEVVVLPEEVAHVLQEARNKGFKT
sp|P09384|CHEA_  LALEAKGETTPAVVETAALSAAIQEESVAETESPRDESKL----------------------

cons             :* ..* *... : . :      .*       . .. *


sp|P07363|CHEA_  ---RIIL---SRLKAGEVDLLEEELGHLTTLTDVVKGADSLSAILPGDIAEDDIT--------
sp|Q56310|CHEA_  FYIKVILKEGTQLKSARIYLVFHKLEELK--CEVVRTIPSVEEIEEE-KFENEVELFVISPVD
sp|P09384|CHEA_  ---RIVL---SRLKANEVDLLEEELGNLATLTDVVKGADSLSATLDGSVAEDDIV--------

cons                :::*   ::**: .: *: .:* .*    :**:   *:.        *:::


sp|P07363|CHEA_  -----AVLCFVIEADQITFETVEVSPK-----------ISTPPVLKLAAEQAP-TGRVEREKT
sp|Q56310|CHEA_  LEKLSEALSSIADIERVI----------------------IKEVTAVTEESGA-EKRTEKEEK
sp|P09384|CHEA_  -----AVLCFVIEADQIAFEKVVAAPVEKAQEKTEVAPVAPPAVVAPAAKSAAHEHHAGREKP

cons                  .*. : : :::                         *   : :...   :. :*:


sp|P07363|CHEA_  --------TRS-NESTSIRVAVEKVDQLINLVGELVITQSMLAQRSSELDPVNHGDLITSMGQ
sp|Q56310|CHEA_  TEKTEEKAERKKVISQTVRVDIEKLDNLMDLMGELVIARSRILE---TLKKYNIKELDESLSH
sp|P09384|CHEA_  --------ARE-RESTSIRVAVEKVDQLINLVGELVITQSMLAQRSNELDPVNHGDLITSMGQ
```

```
cons                      *.   * ::** :**:*:*::*:*****::* : :    *.  *  :*  *:.:

                                 -------α1--------             ------α2--
sp|P07363|CHEA_     LQRNARDLQESVMSIRMMPMEYVFSRYPRLVRDLAGKLGKQVELTLVGSSTELDKSLIERIID
sp|Q56310|CHEA_     LSRITLDLQNVVMKIRMVPISFVFNRFPRMVRDLAKKMNKEVNFIMRGEDTELDRTFVEEIGE
sp|P09384|CHEA_     LQRNARDLQESVMSIRMMPMEYVFSRFPRLVRDLAGKLGKQVELTLVGSSTELDKSLIERIID

cons                *.* : ***: **.***:*:.:**.*:**:***** *:.*:*:: : *..****::::*.* :

                    ----α2------     ---α3--                           -----α4--
sp|P07363|CHEA_     PLTHLVRNSLDHGIELPEKRLAAGKNSVGNLILSAEHQGGNICIEVTDDGAGLNRERILAKAA
sp|Q56310|CHEA_     PLLHLLRNAIDHGIEPKEERIAKGKPPIGTLILSARHEGNNVVIEVEDDGRGIDKEKIIRKAI
sp|P09384|CHEA_     PLTHLVRNSLDHGIEMPEKRLEAGKNVVGNLILSAEHQGGNICIEVTDDGAGLNRERILAKAM

cons                ** **:**::*****  *:*:  **  :*.*****.*:*.*: *** *** *:::*:*: **

                    ---           --α5-                         --α6--
sp|P07363|CHEA_     SQGLTV---SENMSDDEVAMLIFAPGFSTAEQVTDVSGRGVGMDVVKRNIQKMGGHVEIQSKQ
sp|Q56310|CHEA_     EKGLIDESKAATLSDQEILNFLFVPGFSTKEKVSEVSGRGVGMDVVKNVVESLNGSISIESEK
sp|P09384|CHEA_     SQGMAV---NENMTDDEVGMLIFAPGFSTAEQVTDVSGRGVGMDVVKRNIQEMGGHVEIQSKQ

cons                .:*:       .::*:*:  ::*.***** *:*::************. ::.:.* :.*:*::


sp|P07363|CHEA_     GTGTTIRILLPLTLAILDGMSVRVADEVFILPLNAVMESLQPREADLHPLAGGERVLEVRGEY
sp|Q56310|CHEA_     DKGTKVTIRLPLTLAIIQALLVKVNNLVYAIPIANIDTILSISKEDIQRVQD-RDVIVIRGEV
sp|P09384|CHEA_     GSGTTIRILLPLTLAILDGMSVRVAGEVFILPLNAVMESLQPREEDLHPLAGGERVLEVRGEY

cons                ..**.: * *******::.: *:* . *: :*:  :   *.  : *:: : . . *: :***


sp|P07363|CHEA_     LPIVELWKVFNVAGAKTEATQGIVVILQSGGRRYALLVDQLIGQHQVVVKNLESNYRKVPGIS
sp|Q56310|CHEA_     IPVYRLWEVLQIEH-KEELEEMEAVIVRVGNRKYGIVVDDLLGQDDIVIKSLGKVFSEVKEFS
sp|P09384|CHEA_     LPLVELWKVFDVDGAKTEATQGIVVILQSAGRRYALLVDQLIGQHQVVVKNLESNYRKVPGIS

cons                :*: .**:*:::   * *  :  .**:: ..*:*.::**:*:**.::*:*.* . : :*  :*


sp|P07363|CHEA_     AATILGDGSVALIVDVSALQAINREQRMANTAA
sp|Q56310|CHEA_     GAAILGDGSIALIINVSGIV-------------
sp|P09384|CHEA_     AATILGDGSVALIVDVSALQGLNREQRMAITAA

cons                .*:******:***::**.:
```

**Table S3 Percentage of P1 dimer in the "upper" and "lower" density regions in the ATP-free and ATP-bound systems.** A COM distance cutoff of 25 Å was used to account for possible P1 dimer configurations. The "Overall" column reports the P1 dimer percentage in all configurations from all combined simulation trajectories for both systems.

| systems | "upper" | "lower" | Overall |
|---|---|---|---|
| ATP-free | 1.43% | 35.81% | 11.29% |
| ATP-bound | 0.56% | 29.46% | 13.41% |

### CHARMM topology and parameter files of HyRes ATP

```
*topology file for coarse grained RNA
*
   20    1
MASS   400 AP    78.971     ! 1-3; mapping scheme
MASS   401 AO1   30.026     ! 4
MASS   402 AO2   42.037     ! 5
MASS   403 AO3   153.012    ! 6
MASS   404 AN1   39.037     ! 7
MASS   405 AN2   26.018     ! 8
MASS   406 AN3   27.026     ! 9
MASS   407 AN4   42.041     ! 10


RESI ATP         -4.00  !
GROUP                   !
ATOM P3   AP     -2.00  !
ATOM P2   AP     -1.00  !                  (8)  NB—ND  (10)
ATOM P1   AP     -1.00  !                       |  |
GROUP                   !                  (7)  NA—NC  (9)
ATOM C1   AO1     0.00  !     (1) (2) (3)  (4)  |
ATOM C2   AO2     0.00  !     P3--P2--P1---C1---C2 (5)
ATOM C3   AO3     0.00  !                       |
GROUP                   !                       C3 (6)
ATOM NA   AN1     0.00  !
ATOM NB   AN2     0.00  !
ATOM NC   AN3     0.00  !
ATOM ND   AN4     0.00  !
BOND P3   P2        P2   P1       P1   C1       C1   C2
BOND C2   C3        C2   NA       NA   NB       NA   NC
BOND NB   ND        ND   NC       NB   NC
ANGL P3   P2   P1        P2   P1   C1       P1   C1   C2
ANGL C1   C2   C3        C1   C2   NA       C3   C2   NA
ANGL C2   NA   NB        C2   NA   NC
IMPR ND   NC   NB   NA        ND   NC   NA   NB
IMPR ND   NB   NA   NC        NC   ND   NB   NA        NB   ND   NC   NA

END
```

```
BOND
!
!V(bond) = Kb(b - b0)**2
!
!Kb: kcal/mole/A**2
!b0: A
!
!atom type Kb          b0
!
! force constant / 418.4 --> kcal / mol / A^2
AP    AP    35.85       3.00 ! 1-2 bond: length: 0.3 nm; force constant: 30000 kj / mol / nm^2  (in itp)
AP    AO1   100.0       2.25 ! 3-4 bond: length: 0.225 nm; constraint  (in itp)
AO1   AO2   45.85       2.90 ! 4-5 bond: length: 0.29 nm; force constant: 15000 kj / mol / nm^2  (in itp)
AO2   AO3   100.0       2.25 ! 5-6 bond: length: 0.225 nm; constraint  (in itp)
AO2   AN1   59.75       2.28 ! 5-7 bond: length: 0.288 nm; force constant: 50000 kj / mol / nm^2  (in itp)
AN1   AN2   100.0       1.60 ! 7-8 bond: length: 0.160 nm; constraint
AN1   AN3   100.0       2.81 ! 7-9 bond: length: 0.281 nm; constraint
AN2   AN4   100.0       2.45 ! 8-10 bond: length: 0.245 nm; constraint
AN3   AN4   100.0       2.62 ! 9-10 bond: length: 0.262 nm; constraint
AN2   AN3   100.0       3.08 ! 8-9 diagonal distance; constraint

THETAS
! angles in ATP
AP  AP  AP  6.0     127.5
AP  AP  AO1 4.0    87.7
AO1 AO2 AO3 10.0    83.0
AP  AO1 AO2 15.0   138.1
AN1 AO2 AO3 25.0   96.0
AO1 AO2 AN1 4.5    145.7
AO2 AN1 AN2 42.5   162.0
AO2 AN1 AN3 5.75    114.0


IMPHI
! 70/4.184 = 16.73 *2
AN1 AN2 AN3 AN4 33.46   0   180.0
AN2 AN1 AN3 AN4 21.51   0   0.0
AN3 AN1 AN2 AN4 43.02   0   0.4
AN1 AN2 AN4 AN3 43.02   0   -0.4
AN1 AN3 AN4 AN2 43.02   0   0.4
NONBONDED  NBXMOD 3  ATOM CDIEL SWITCH VATOM VDISTANCE VSWITCH -
     CUTNB 16.0  CTOFNB 15.0  CTONNB 11.  EPS 1.0  E14FAC 0.4  WMIN 1.5

! ATP
AP       0.00     -0.15042      2.214    0.00     -0.15042      2.214
AO1      0.00     -0.12033      2.000    0.00     -0.12033      2.000
AO2      0.00     -0.12033      2.274    0.00     -0.12033      2.274
AO3      0.00     -0.13538      2.417    0.00     -0.13538      2.417
AN1      0.00     -0.10153      2.261    0.00     -0.10153      2.261
AN2      0.00     -0.09025      2.000    0.00     -0.09025      2.000
AN3      0.00     -0.09025      2.000    0.00     -0.09025      2.000
AN4      0.00     -0.10153      2.239    0.00     -0.10153      2.239

HBOND AEXP 10 REXP 12 HAEX 4 AAEX 2   NOACCEPTORS  HBNOEXCLUSIONS  ALL  -
   CUTHB 6.0 CTOFHB 5.0 CTONHB 4.0  CUTHA 100.0  CTOFHA 90.0  CTONHA 90.0


END
```
